# Modeling the transmission dynamics of African swine fever virus within commercial swine barns: Quantifying the contribution of multiple transmission pathways

**DOI:** 10.1101/2024.08.02.604486

**Authors:** Aniruddha Deka, Jason A Galvis, Christian Fleming, Maryam Safari, Chi-An Yeh, Gustavo Machado

## Abstract

Transmission of the African swine fever virus (ASFV) within commercial swine barns occurs through direct and indirect pathways. Identifying and quantifying the roles of ASFV dissemination within barns is crucial for the development of effective disease control strategies. We developed a stochastic transmission model to examine the ASFV dissemination dynamics through eight transmission routes within commercial swine barns. We consider seven transmission routes at three disease dynamics levels: within-pens, between-pens, and within-room transmission, along with the transfer of pigs between pens within the same room. We simulated ASFV spread within barns of various sizes and layouts from rooms with a median of 32 pens (IQR:28-40), where each pen housing a median of 34 pigs (IQR: 29-36). Our model enables the tracking of the viral load in each pen and the monitoring of the disease status at the pen level. Simulation results show that between-pen transmission pathways exhibited the highest contribution to ASFV spread, accounting for 71.4%, where within-pen and within-room pathways account for 20.1% and 8.5%, respectively. Among the direct transmission pathways, nose-to-nose contact between pens was the primary route of dissemination, comprising an average of 49%, while the fecal transmission between pens contributed 21%. On the other hand, aerosol transmission within pens had the lowest contribution, accounting for less than 1%. Furthermore, we show that the daily transfer of pigs between pens did not impact the spread of ASFV. The combination of passive surveillance of daily detection and active surveillance focused on mortality allowed the detection of ASFV within three Days, with peak detection occurring when mortality rates peaked. The model also allows us to pinpoint where the majority of infections and viral load are concentrated during the ASFV spread. This work significantly deepens our understanding of ASFV spread within commercial swine production farms in the U.S. and highlights the main transmission pathways that should be prioritized when implementing ASFV countermeasure actions at the room level.

## 1. Introduction

The swine industry is vulnerable to infectious diseases that cause substantial production losses, especially foreign animal diseases such as the African swine fever virus (ASFV) [1–3]. After its initial detection in Portugal in 1957, ASFV spread throughout Western Europe over the subsequent 30 years. It later surfaced in Eastern Europe in 2007, rapidly affecting neighboring countries [4–6]. ASFV was identified in Asia in 2018, where it continues to spread [7, 8]. Once introduced into commercial swine farms, ASFV dissemination within pig barns is facilitated by multiple transmission pathways (e.g., direct contact, environment), making it imperative to comprehend the contributions of such pathways in developing and refining within-farm mitigating strategies [9, 10]. Numerous studies have evaluated the probability of ASFV introduction and propagation among farms [11–15]. Understanding the complexity of ASFV transmission within densely populated facilities, which for commercial swine production are barns that are often subdivided into rooms and further into pens, remains to be examined [16]. An alternative solution to the lack of fieldwork data on ASFV dynamics within barns is the development of disease transmission models [17–19]. Most ASFV transmission models have been at the farm level and have explored various dissemination pathways of ASFV, including domestic-to-domestic transmission and wild boar-to-domestic [13, 18, 20, 21]. However, limited efforts in simulating ASFV transmission within farms have simplified transmission pathways by focusing on one or two transmission pathways and do not assess the contributions of individual transmission pathways within farm [22, 23].

Upon the introduction of ASFV into commercial swine premises, dissemination is driven by direct contact between infectious and Susceptible animals [24–26]. In commercial swine operations, growing pigs are housed in barns that are further subdivided into pens. For instance, a room inside a barn is subdivided into pens by open bar panels made of metal bars that allow the arrangement of pen space as needed. However, it facilitates direct interaction with neighboring pigs, potentially leading to between-pen transmission [16]. The indirect transmission process entails infectious pigs shedding the virus through bodily excretions, and secretions in feces and exposing Susceptible animals [27–30]. Such infectious materials can also become aerosolized when pigs sneeze or cough, thus posing a risk for short-distance spread within a barn room [31]. Furthermore, once dried, these secretions and excretions have been found in dust, which may also be aerosolized and potentially lead to aerosol transmission [31–35]. In addition, human activities and the limited biosecurity measures have significantly contributed to the spread of ASFV via contaminated materials equipment or contaminated boots among other fomites [36, 37]. Further, the practice of sorting pigs (moving pigs between pens) is typically aimed at improving growth, health, and slaughter readiness [38, 39]. However, this approach can inadvertently transmit diseases to previously disease-free pens due to the introduction of sick animals on uninfected pens [38].

Examining the interactions among pigs at different levels within a room is crucial in understanding the transmission dynamics of infectious diseases. Mathematical models that incorporate the dynamics of disease at multiple levels could serve as valuable tools for gaining a deeper understanding of how ASFV may spread. Here, we aim to fill these gaps by modeling ASFV transmission within a significant number of commercial swine barns in operation in the U.S. Thus, we developed an epidemiological model to simulate the within-barn ASFV dissemination dynamics, incorporating the variations of of barn sizes and varying pig capacity in each pens. Our compartmental model considers Susceptible, Exposed, Clinical, SubClinical, Chronic-Carrier, Detected, and Recovered dynamics within-pen, between-pen, and room level. Our model outcome includes the contribution of seven direct within-barn transmission pathways and the role of moving pigs between pens as the indirect transmission route over a period of 175 Days. We evaluate the infectiousness of each pen throughout the ASFV outbreak, providing insights into the intensity and dissemination within barns. Our mathematical model tracks the viral load in pens resulting from fecal and airborne transmission, thereby improving our understanding of disease progression within commercial swine operations. In this study, we investigated two disease surveillance methods: passive surveillance, which involves daily detection of a fixed percentage of Infected pigs, and active surveillance triggered by observed mortality. Both surveillance methods for ASFV-Infected pigs are essential for developing potential intervention strategies should ASFV emerge in the U.S. [23, 40]. However, our model focuses on highly virulent strains to analyze the spread of ASFV. Previous studies have shown that ASFV has been characterized as an extremely contagious disease with mortality rates reaching up to 100% [12, 41, 42]. Yet, various experimental and field studies suggest that the transmission of ASFV is not always rapid [43]. Thus, we assessed the simulated output and infectiousness level across scenarios that included a) human-mediated transmission parameter, b) increased mortality rates for various compartments, and c) varying probability rates for daily pig transfers.

## 2. Methods

### 2.1. Data

This study used data from 1958 commercial swine farms managed by 33 swine production companies in the U.S. To access the different barn layouts of the different farms we have in our database, we used farm maps from enhanced on-farm Secure Pork Supply (SPS) biosecurity plans available in the Rapid Access Biosecurity application(RABapp^™^)[44]. Briefly, The RABapp^™^ results from a multi-state consortium of academic researchers, animal health officials, and swine businesses that serves as a platform for standardizing the approval of SPS biosecurity plans while storing and analyzing animal movement data [44]. SPS biosecurity plans are part of a United States Department of Agriculture (USDA) and Pork Checkoff initiative ^1^ to enhance business continuity by helping swine producers in implementing biosecurity measures on individual farms. The SPS biosecurity plan comprises a written site-specific enhanced biosecurity plan and a farm map composed of fourteen features, including the line of separation (LOS)(Figure 1). Each premises map has at least one LOS established as a control boundary to prevent pathogen movement into areas where Susceptible animals are housed (a.k.a. barns). In RABapp^™^, premises maps are stored as geospatial data. The LOS feature polygons in this data were used to create polygons representing each barn, which allowed us to calculate the barn size and subsequently estimate the number of rooms and pens in each barn; a schematic of the barn is illustrated in Figure 1. Our database had 7317 barns with a total of 7704 rooms, out of which 6930 had 1 room and 387 barns had 2 rooms. With consistent pen sizes used across the industry (the average dimensions of pens are 19.6 ft × 9.8 ft with 15-35 pigs per pen [45], we developed an algorithm that takes as input the area of a pen to generate pens distributed within each room. Out of the 7704 rooms in the database, the median number of pens per room was 32 (interquartile range (IQR): 28-40). The number of pigs in each farm is part of the SPS plans and stored in the RABapp^™^ [44]; this data was used to distribute pigs per pen, which was done by taking the total number of pigs per farm and distributed over the total number of pens. We have detailed the above methodology in the Supplementary material (Section 1.1).

**Fig. 1:**
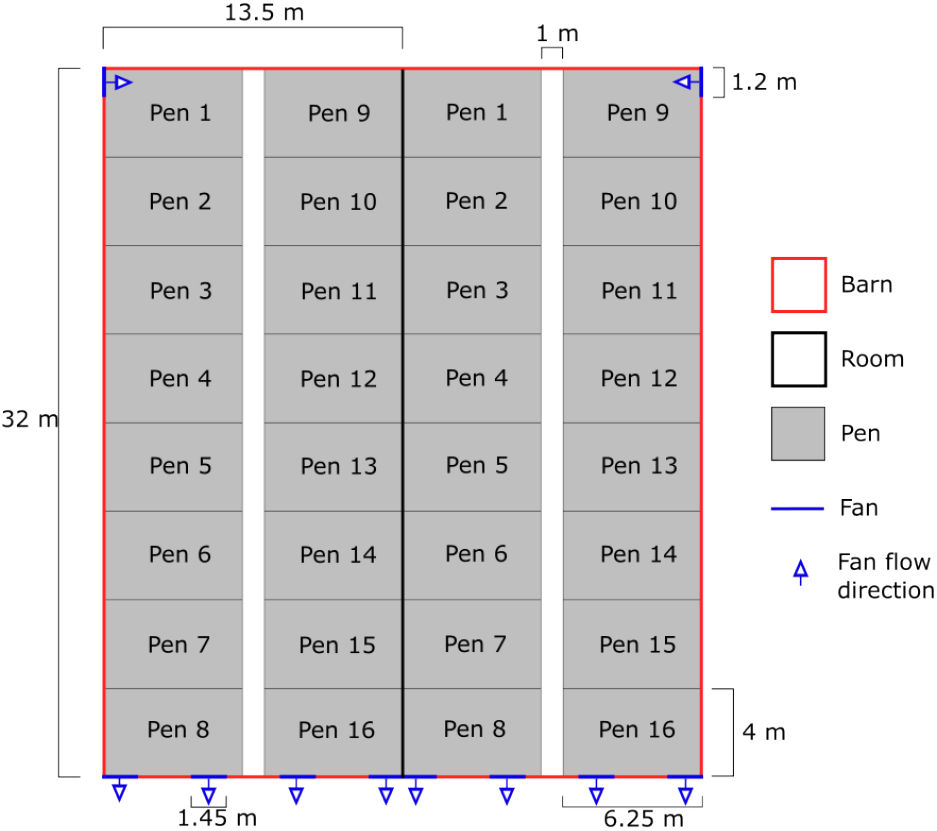
A schematic of a barn with two rooms separated by solid black lines. In this example, each room fitted 16 pens, arranged in two rows of eight pens each. The solid white line represents the walkway between the two rows of pens. The dimension of each pen is defined in the figure.

### 2.2. Within-barn transmission dynamics

Our barn-level pig population is divided into seven health statuses, and the infection pathway for pigs is described in the model as follows: Susceptible (*S*) negative pigs; Exposed (*E*) pigs that have been Exposed to the virus but are not Infected yet; Sub-Clinical (*S* _*c*_) pigs that have been Exposed to the virus but have not yet shown Clinical signs but shed virus into the environment; Clinical (*C*) pigs that have been Exposed and exhibit Clinical signs and shed the virus into the environment; Chronic Carrier (*C*_*c*_) pigs that have been Exposed to the virus but do not show Clinical signs and do not shed the virus into the environment; Detected (*D*) pigs that are either *S* _*c*_, *C* or *C*_*c*_ and that have been successfully Detected; and Recovered (*R*) pigs that were Infected but have fully recovered (Figure 2)

**Fig. 2:**
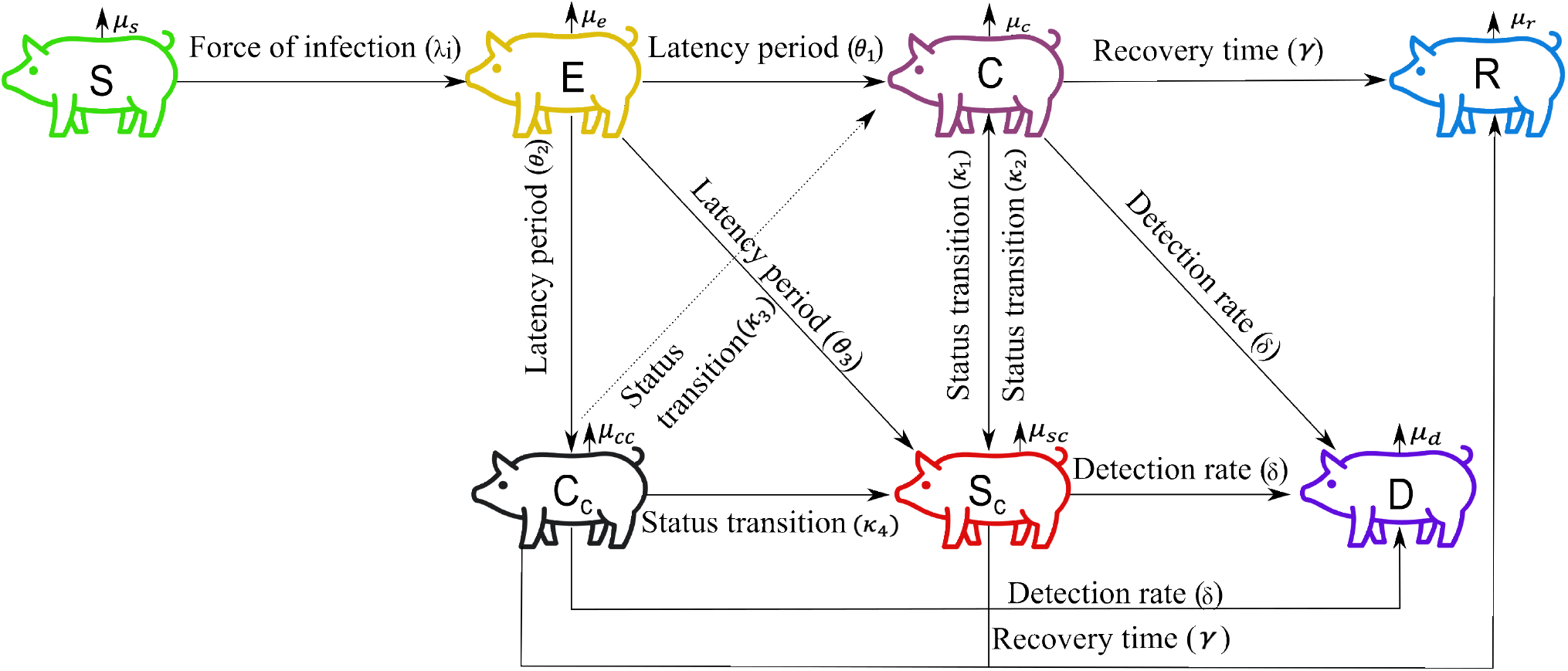
A schematic illustrating the transmission and population dynamics. This diagram outlines the progression of pigs through seven compartments: Susceptible (*S*), Exposed (*E*), Clinical (*C*), Sub-Clinical (*S* _*c*_), Carrier (*C*_*c*_), Detected (*D*), and Recovered (*R*). *λ*_*i*_ represents the force of infection, indicating the rate at which pigs transition from the Susceptible to the Exposed compartment. *i* = 1, 2, …, 7 defines the various pathways through which exposure can occur. *θ* = 1, 2, 3, specifies the transitions of Exposed pigs to Clinical, Sub-Clinical, or Carrier compartments after the latency period. *κ* describes the status changes of pigs within the Clinical, Sub-Clinical, and Carrier compartments. *δ* is the detection rate of Infected pigs, encompassing those in the Clinical, Sub-Clinical, and Carrier compartments. *γ* denotes the recovery rate for pigs in each compartment. Parameters are detailed in full in Table 1 and transition dynamics is defined in Table 2

The within-barn transmission model considers the spread of ASFV at three levels: 1) within pens, 2) between pens, and 3) at room level, allowing ASFV to spread across pens. The model incorporates eight distinct dissemination pathways: two direct routes, i) nose-to-nose contact within pens and ii) noseto-nose contact of pigs between adjacent pens, and six indirect pathways, iii) fecal transmission within pens, iv) air transmission within pens, v) fecal transmission between adjacent pens, vi) room air flow transmission between pens within the room, vii) human-mediated transmission, and viii) movements of pigs between pens.

**Table 1:** Summary of parameters used for the dissemination of African swine fever virus within commercial swine barns.

| Parameter | Meaning | Value | Range/Distribution | Source |
| --- | --- | --- | --- | --- |
| $\beta_{wp}$ | Within-pen direct transmission rate | min = 1.57, mod = 2.79, max = 4.95 | PERT | [57] |
| $\beta_{bp}$ | Between-pen direct transmission rate | min = 0.2, mod = 0.3, max = 0.5 | PERT | [57] |
| $\omega_v$ | Particles shed in faeces post-inf. time | min = $10^4$, max = $4.96 \times 10^6$ /g | Uniform | [51] |
| $V_{shed}$ | Particles spread through aerosol | min = $10^2$, max = $10^{3.2} TCID_{50} eq./m^3$ | Fixed | [35] |
| $V_{area}$ | Dimension of pen | $25.85 m^3$ | Fixed | Calculated |
| $\beta_{fp}$ | Fecal transmission rate within and between pen | $10^{-2}$ | Fixed | [54] |
| $\beta_{ap}$ | Aerosol transmission rate | min = $6 \times 10^{-3}$, $6 \times 10^{-2}$, max = $9 \times 10^{-2}$ | PERT | [35] |
| $\epsilon_1$ | Rate at which the virus decays | 0.08 | Fixed | [53] |
| $\epsilon_2$ | Fecal removal per g/per Day (Fraction) | 0.7 | Fixed | [53] |
| $Q_i$ | Fecal ingestion rate by finisher pigs | 25 (g/Day) | Fixed | [88] |
| $Q_{shed}$ | Average faeces shed by a finisher pig | 1000 (g/Day) | Fixed | [89] |
| $C_\beta$ | Human transmission rate | 0.01 | 0-1 | [90] |
| $1/\theta$ | Length of latency period (Days) | min = 6, max = 10 | Uniform | [20] |
| $1/\gamma$ | Length of infectious period (Days) | min = 10, max = 38 | Uniform | [20] |
| $1/\mu_s$ | Average mortality rate of Susceptible | 0.01 | Fixed | [65] |
| $1/\mu_e$ | Average mortality rate of Exposed | 0.05 | Fixed | Assumed |
| $1/\mu_c$ | Average mortality rate of Clinical | 0.06 | Fixed | [65] |
| $1/\mu_{sc}$ | Average mortality rate of Sub-Clinical | 0.06 | Fixed | [65] |
| $1/\mu_{cc}$ | Average mortality rate of Chronic-Carrier | 0.01 | Fixed | Assumed |
| $1/\mu_d$ | Average mortality rate of Detected | 0.06 | Fixed | [65] |
| $\kappa_1$ | Transition rate from Clinical to Sub-Clinical | min = 0, max = 0.01 | Uniform | Assumed |
| $\kappa_2$ | Transition rate from Sub-Clinical to Clinical | min = 0, max = 0.1 | Uniform | Assumed |
| $\kappa_3$ | Transition rate from Chronic-Carrier to Clinical | min = 0, max = 0.01 | Uniform | Assumed |
| $\kappa_4$ | Transition from Chronic-Carrier to Sub-Clinical | min = 0, max = 0.01 | Uniform | Assumed |
| $\theta_1$ | Exposed pigs that become Clinical (Fraction) | 0.8 | 0-1 | Assumed |
| $\theta_2$ | Exposed pigs that become Sub-Clinical (Fraction) | 0.05 | 0-1 | Assumed |
| $\theta_3$ | Exposed pigs that become Chronic-Carrier (Fraction) | 0.15 | 0-1 | Assumed |
| $\nu$ | Daily detection rate of Infected | 0.01 | 0-1 | Assumed |
| $P_{dh1}$ | Detection rate for 10% mortality (Clinical) | 0.95 | 0-1 | Assumed |
| $P_{dh2}$ | Detection rate for 10% mortality (Sub-Clinical) | 0.75 | 0-1 | Assumed |
| $P_{dh3}$ | Detection rate for 10% mortality (Chronic-Carrier) | 0.5 | 0-1 | Assumed |
| $P_{dl1}, P_{dl2}, P_{dl3}$ | Detection rate for low mortality | 0.01 | 0-1 | Assumed |
| $T_{sen}$ | Test-sensitivity (Fraction) | 0.9 | 0-1 | Assumed |
| $P_{ij}$ | Pig transfer probability per Day | 0.05 | 0-1 | Assumed |
| $\eta_1$ | Fraction of fecal shed within pen | 0.9 | 0-1 | Assumed |
| $\eta_2$ | Fraction of fecal shed between pens | 0.1 | 0-1 | Assumed |
| $\xi_1$ | Fraction of fecal ingested within pen | 0.9 | 0-1 | Assumed |
| $\xi_2$ | Fraction of fecal ingested between pens | 0.1 | 0-1 | Assumed |

**Table 2:** Transition dynamics between compartments and their occurrence rates.

| Status | E | C | $S_c$ | $C_c$ | $D$ | $R$ |
| --- | --- | --- | --- | --- | --- | --- |
| $S$ | $\lambda S$ | × | × | × | × | × |
| $E$ | × | $\theta_1 E$ | $\theta_2 E$ | $\theta_3 E$ | × | × |
| $C$ | × | × | $\kappa_1 C$ | × | $(\nu + P_{dl1} \times P(T_{sen}))C, \mu < 10\%$<br>$(\nu + P_{dh1} \times P(T_{sen}))C, \mu \geq 10\%$ | $\gamma C$ |
| $S_c$ | × | $\kappa_2 S_c$ | × | × | $(\nu + P_{dl2} \times P(T_{sen}))S_c, \mu < 10\%$<br>$(\nu + P_{dh2} \times P(T_{sen}))S_c, \mu \geq 10\%$ | $\gamma S_c$ |
| $C_c$ | × | $\kappa_3 C_c$ | $\kappa_4 C_c$ | × | $(\nu + P_{dl3} \times P(T_{sen}))C_c, \mu < 10\%$<br>$(\nu + P_{dh3} \times P(T_{sen}))C_c, \mu \geq 10\%$ | $\gamma C_c$ |
| $D$ | × | × | × | × | × | $\gamma D$ |

Here, $\lambda$ denotes the rate of exposure through the transmission pathways outlined in the main text. $\theta_1, \theta_2, \theta_3$ define the length of the latency period. $\kappa_1, \kappa_2, \kappa_3, \kappa_4$ define the transition rates from Clinical to Sub-Clinical, Sub-Clinical to Clinical, Chronic-Carrier to Clinical, and Chronic-Carrier to Sub-Clinical respectively. $\nu$ is the daily detection rate of infected pigs. $P_{dh1}, P_{dh2}, P_{dh3}$ define the detection rates for a 10% mortality rate for Clinical, Sub-Clinical, and Chronic-Carrier pigs, respectively. $P_{dl}$ is the detection rate for a low mortality rate. $\mu$ is the mortality rate of pigs, and $\gamma$ is the recovery rate of pigs.

We assumed the initial population is Susceptible (*S*), given that the U.S. remains ASFV-free. The probability of infection governs the transition from the *S* compartment to the *E* compartment, denoted as *λ*_*i*_ for seven transmission routes (*i* = 1, 2, …, 7) (Supplementary Material Figures S1-S7) and additionally due to the transfer of pigs between pens. The number of pigs in the *E* compartment is influenced by the latency period (*θ*). Latency is drawn from a uniform distribution with a minimum of 6 Days and a maximum of 10 Days (Table 1). Exposed pigs, after the latency period, follow one of three paths: a) Clinical (*C*) pigs shed the virus into the environment and spread the disease, b) Sub-Clinical (*S* _*c*_) pigs also shed the virus and contribute to the spread, and c) Chronic-Carrier (*C*_*c*_) pigs have been Exposed to the virus but do not show Clinical signs nor shed the virus into the environment (Figure 2). Our model assumes that following an exposure event, the majority of pigs become Clinical, a small fraction become Sub-Clinical, and very few become chronic Carriers (Table 1). We consider 0 ≤ *θ*_1_, *θ*_2_, *θ*_3_ ≤ 1, *θ*_1_ + *θ*_2_ + *θ*_3_ = 1 (Table 1) where *θ*_1_, *θ*_2_, *θ*_3_ represents the fraction of the Exposed population that, after the latency period, becomes Clinical, Sub-Clinical and Chronic-Carrier, respectively. Furthermore, our model accounts for Clinical pigs transitioning to the Sub-Clinical state and vice versa at the rate of *κ*_1_&*κ*_2_ (0 *< κ*_1_, *κ*_2_ < 1), respectively. Moreover, Chronic-Carriers can transition to Clinical and SubClinical status at the rate of *κ*_3_ and *κ*_4_ (0 < *κ*_3_, *κ*_4_ < 1), respectively, whereas Clinical and Sub-Clinical individuals do not transition to Chronic-Carriers. We also consider mortality among all status, albeit at a different rate (*µ*) with maximum mortality in the Clinical (*µ*_*c*_), Sub-Clinical (*µ*_*sc*_) and Detected compartments (*µ*_*d*_), followed by, Exposed (*µ*_*e*_), Susceptible (*µ*_*s*_), and Chronic-Carrier compartments (*µ*_*cc*_). The mortality rate for each compartment is defined in Table 1. Given that we do not implement control actions, Detected pigs are not removed from our simulation system, and thereby, they contribute to disease transmission. An infectious pig recovers at a rate *γ*, which is simulated as a uniform distribution in our simulations (Table 1). It is important to highlight that our model does not consider reinfection; therefore, recovered pigs maintain their disease-free status without possibly reacquiring the infection. The transition among compartments and the corresponding rates of these transitions are illustrated in Table 2. To account for disease detection (*µ*_*δ*_), we consider two surveillance rates: a) passive surveillance, where a fixed number of Infected pigs (1% of Infected pigs comprising Clinical, Sub-Clinical and Carrier pigs) are Detected each Day to estimate ASFV spread, and b) active surveillance, triggered by mortality rates. It is important to note that the mortality threshold for activating surveillance is assessed individually for each pen. For passive surveillance, Clinical, Sub-Clinical, and Chronic-Carrier pigs are Detected at a specific rate per Day (Table 1); b) active surveillance, on the other hand, is initiated based on a predefined mortality rate threshold *µ*_*δ*_, where (*µ*_*δ*_ = *µ*_*s*_ + *µ*_*e*_ + *µ*_*c*_ + *µ*_*sc*_ + *µ*_*cc*_ + *µ*_*d*_ + *µ*_*r*_), representing deaths of Susceptible, Exposed, Clinical, Sub-Clinical, Carrier, Detected and recovered pigs respectively. If mortality surpasses the mortality threshold *µ*_*δ*_, the detection rate is adjusted as detailed in Equation 2. It should be noted that in passive surveillance, all Infected pigs (Clinical, Sub-Clinical, and chronic-Carrier) have an equal probability of being Detected. In contrast, active surveillance has varying detection probabilities: 80% for Clinical pigs, 15% for Sub-Clinical pigs, and 5% for chronic-Carrier pigs. The percentage of pigs Detected through mortality-based surveillance in each pen can be defined as follows

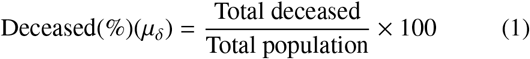

We also consider diagnostic test sensitivity of 90% as a conservative value due to the variations among commercial kits [46].

The total number of Detected pigs due to active surveillance can be expressed as follows

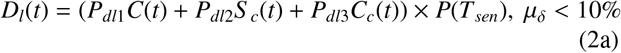

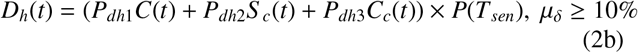

Where *D*_*l*_(*t*) and *D*_*h*_(*t*) are the Detected pigs for low and high rates of mortality, respectively. *P*_*dl*1_, *P*_*dl*2_, *P*_*dl*3_ are the detection rate for Clinical, Sub-Clinical and Carrier pigs, respectively with *P*_*dl*1_ = *P*_*dl*2_ = *P*_*dl*3_, for low mortality rate, and the detection rates of *P*_*dh*1_, *P*_*dh*2_ and *P*_*dh*3_ are corresponding to Clinical, Sub-Clinical and Carrier pigs, respectively, for high mortality rate.

We also monitor the pen status in each room daily by the following criteria: 1) pen is deemed disease-free on a Day if all pigs within the pen are Susceptible; 2) pen is considered Infected if at least one pig within the pen is in a Clinical, SubClinical, or Chronic Carrier state and there is no Detected pig; 3) pen is classified as Detected if at least one pig remains Detected within pen.

### 2.3. Within-pen transmission

#### 2.3.1. Within pen nose-to-nose contact

Most swine infectious pathogens, including ASFV, disseminate through direct contact between healthy pigs (*S*) and Infected pigs (*I*), where the latter is represented in our model as Clinical, Sub-Clinical, and Detected pigs [47, 48] (Supplementary Material Figure S1). Here, we assumed that transmission via direct contacts occurring exclusively within a pen is frequencydependent. Furthermore, we do not consider different behaviors within a group of pigs, for example, aggressive behavior and dominance relationships within pigs, and assume a consistent rate of contact among pigs [49, 50]. The within-pen direct contact force of infection is modulated by

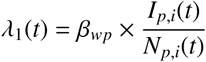

where *β*_*wp*_ modulates within the pen ASFV transmission rate. The Malta variant was chosen to illustrate direct nose-to-nose contact [35] defined in Table 1. *I*_*p,i*_(*t*) and *N*_*p,i*_(*t*) denote the number of Infected and total pigs available in the particular pen per unit of time *t*, respectively. The probability that a Susceptible pig is Exposed to ASFV due to contact with Infected pigs in a pen is given by

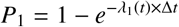

#### 2.3.2. Within pen oro-fecal transmission

Shedding pathogens through feces plays a crucial role in transmitting diseases within swine barns [51]. ASFV-Infected pigs contaminate the environment through excretions and secretions via oral fluid, nasal fluid, feces, and urine, which is exceptionally high during the acute phase [52]. To model these dynamics, we used a similar approach of [53]. We used *ω*_*v*_ to represent the average amount of virus shed by an infectious pig per gram of feces each Day and is defined as a uniform distribution that spans between the minimum and maximum viral loads (Table 1), which is then multiplied by the daily fecal output 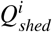 from Infected pigs. We also consider fecal shedding into adjacent pens, which is captured by 0 *< η*_1_, *η*_2_ < 1 (Table1) where *η*_1_, *η*_2_ indicate the fractions of fecal matter shed within the pen and into adjacent pens, respectively, with the condition that *η*_1_ +*η*_2_ = 1. Notably, if a pen has two neighboring pens, the fecal matter that can be introduced into each neighboring pen is *η*_2_*/*2. Additionally, we incorporate *ϵ*_1_ and *ϵ*_2_ for the daily removal of fecal matter through slatted floors and the environmental decay rate of ASFV, respectively. The accumulation of the virus in feces within a pen per Day is calculated as follows,

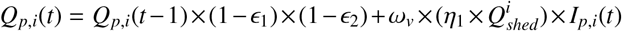

where *Q*_*p,i*_(*t*) denotes the virus concentration in fecal matter at time *t* in pen *i*. The force of infection within the pen, *λ*_2_(*t*), is represented by

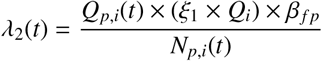

A schematic of the transmission route is described in Supplementary Material Figure S2. The parameter *β*_*fp*_ represents the environmental contamination rate from infectious fecal matter. The Georgia variant was used to investigate fecal transmission [54]. The transmission rate *β*_*fp*_ within a pen, corresponds to the average number of animals that can be Infected by a single viral cell present in the pen environment (*β*_*fp*_ = 10^−2^). This rate inversely relates to the required number of viral particles needed in the environment to infect one pig [53, 55]. In short, the inverse of (*β*_*fp*_ = 10^−2^) corresponds to the average number of viral copy number of viral cells per gram of feces in the environment required to infect one animal in one Day, i.e., 100 cells/g/ Day (Table 1). The dissemination by this route also depends on the ingestion of Infected fecal matter by Susceptible pigs. *Q*_*i*_ is the daily amount of feces ingested by an animal. We consider that pigs ingest a fraction of the fecal denoted as 0 < *ξ*_1_ < 1 within the pen and the remaining 0 < *ξ*_2_ < 1 from its neighboring pens with the condition *ξ*_1_ + *ξ*_2_ = 1. For pens with two adjacent pens (Supplementary Material Figures S2 and S5), the fecal ingested from each neighboring pen is *ξ*_2_*/*2. *N*_*p,i*_(*t*) and *I*_*p,i*_(*t*) denote the total number of pigs and Infected pigs in pen *i* at time *t*, respectively. The probability that a Susceptible pig becomes Exposed to ASFV through oro-fecal transmission is given by:

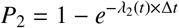

#### 2.3.3. Within pen air-flow transmission

ASFV dissemination via air has been demonstrated especially at short distances (e.g., within a meter) [31, 56, 57]. After the latency period, Clinical, Sub-Clinical, and Detected pigs shed viral particles into the environment, where they remain viable and contribute to the contamination of the environment [51]. We consider that the virus decays at a fixed rate (*ϵ*_1_) within a pen, which is assumed to be equal to the oro-fecal virus decay rate. The shedding rate of aerosol virus per cubic meter of the air by an infectious pig, 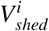, is characterized by a uniform distribution that encompasses the range from the minimum to the maximum viral load shed (Table 1). A schematic of the transmission route is described in Supplementary Material Figure S3. The viral load released at time *t* is given by:

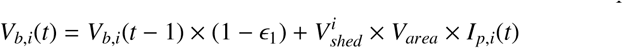

We used our observed pen dimensions (*V*_*area*_) to estimate the total viral load present in the air within each pen (2.1). Thus, the probability that a Susceptible pig becomes Exposed through contaminated airflow within a pen is given by:

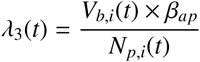

Here *β*_*ap*_ denotes the aerosol transmission rate of ASFV within a pen. We used parameter values from the Brazil variant [35] for aerosol transmission. The probability that Susceptible pigs are Exposed through airflow is given by

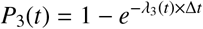

The overall probability of a Susceptible pig getting Exposed within a pen through the three aforementioned transmission routes is defined as

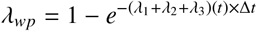

### 2.3. Between-pen transmission

#### 2.4.1. Nose-to-nose pig contact between adjacent pen(s) transmission

We constructed a pen-to-pen network for each room to model dissemination via nose-to-nose contact between pen pigs in adjacent pens[22]. Supplementary Material Figure S4 provides a schematic representation of the transmission pathways. Briefly, rooms within the barn layout feature two longitudinal rows of pens divided by a walkway one meter wide (Figure 1). We defined *G* = (*V, E*) as a graph where *V*_*i*_ = *v*_1_, *v*_2_, …, *v*_*n*_ denotes the set of vertices, with each vertex *v*_*i*_, *i* = 1, 2, …, *n* corresponding to an individual pen within room. *E*_*i*_ is the set of directed edges between vertices, where the edge (*v*_*i*_, *v* _*j*_) represents the adjacency relation between pen *i* and pen *j*. In this pen-to-pen network, connections are defined based on immediate adjacency. For any given pen *i*, there are edges to *v*_*i* −1_ (the left neighbor) and *v*_*i*+1_ (the right neighbor), illustrating the direct neighboring relationships facilitated by metal pen stalls that separate the pens Supplementary Material Figure S4. In evaluating interactions with neighboring pens, we account for a pig’s total potential contacts on any given Day, encompassing those within its pen and with pigs in adjacent pens. We adopt a similar approach to the one discussed in [22] to avoid overestimating daily contact frequency and appropriately parameterize the transmission rate of direct nose-to-nose contact. The likelihood of an Infected pig transmitting the disease to a Susceptible pig in an adjacent pen is thus quantified by

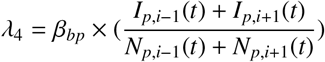

For pens with just one neighboring pen, say *i* + 1 contact rate is divided by two

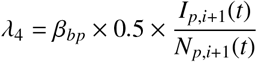

*β*_*bp*_ is the transmission rate of the virus due to direct contact with adjacent pens. The probability of Susceptible pigs getting Exposed through this route is given by

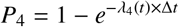

#### 2.4.2. Between pen oro-fecal transmission

Pigs shed feces into their neighboring pens, and therefore, contaminated fecal matter from infectious pigs can be transferred to adjacent pens (Supplementary Material Figure S5). For a pen *i*, the viral load that is accumulated due to the shedding of Infected fecal matter from adjacent pens can be defined as:

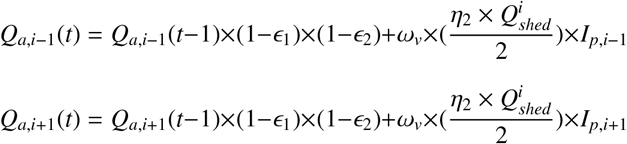

where *Q*_*a,i* −1_ and *Q*_*a,i*+1_ represent the viral load in the feces present at time *t* adjacent to pen *i*, and 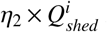 is the fraction of fecal materials shedded in pen *i* from neighboring pens. Note that *I*_*p,i* −1_(*t*) and *I*_*p,i*+1_(*t*) indicate the number of Infected pigs in the neighboring pens, whereas *N*_*p,i* −1_(*t*) and *N*_*p,i*+1_(*t*) denote the total pig population in those pens. The following equation quantifies the likelihood of pigs in pen *i* getting Infected due to ingestion of contaminated fecal matter from adjacent pens:

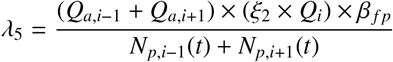

where *λ*_5_ quantifies the probability of exposure in a specific pen *i* resulting from the dispersal of infectious fecal matter from Infected pigs to adjacent pens. *Q*_*i*_ represents the daily average quantity of feces ingested by pigs, and *ξ*_2_ is the proportion of fecal matter that pigs in pen *i* ingest from adjacent pens. *β*_*fp*_ is the rate of environmental transmission as outlined in section 2.3.2. The likelihood of a Susceptible pig in pen *i* becoming Exposed due to ingesting contaminated feces from neighboring pens is as follows:

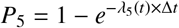

The overall probability of a Susceptible pig getting Exposed through the between-pen transmission routes can be defined as

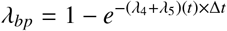

#### 2.4.3. Room-level air-borne transmission

We consider a network in which airflow disseminates infectious viral particles between pens at the room level, in which any animal was likely to be Exposed. We used computational fluid dynamics (CFD) to model the dissemination of infectious viral particles between pens within the room by modeling the mechanical ventilation of the room. This approach extends the algorithm previously developed and published in our earlier work, which is detailed further in [10]. For this route of transmission, a pen-to-pen network is constructed through a directed graph as *G* = (*V, E*), where pens are defined as a set of vertices *V* = *v*_1_, *v*_2_, …, *v*_*n*_ and their directed edges, *E*, signifies the potential pathway between pens, where the edge (*v*_*i*_, *v* _*j*_) determines the amount of airborne pathogens propagated from the pen *j* to pen *i*. In this airborne transmission route, each pen *v*_*i*_ is connected to every other pen *v* _*j*_ where *i* ≠ *j*. As shown in (Supplementary Material Figure S6), edges *E* include all potential pathways between pens (*v*_*i*_, *v* _*j*_) where *i* ≠ *j*, which indicates that the within-pen airborne transmission, *i* = *j*, is excluded, as this route of transmission is considered in Section 2.3.3. Hence, the directed graph, *G*, is defined as,

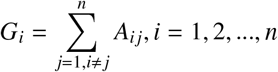

where the *ij* entry of the transmission matrix *A* determines the propagated virus to pen *i* from pen *j*, and *n* is the number of pens in each room. To model the virus transportation to adjacent pens, we assess the generated viral load in each pen based on the number of Infected animals present at each time step within that specific pen. The viral load depends upon the number of particles shed by an infectious pig per Day defined by the parameter 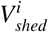 per cubic meter of the air. The level of viruses that are generated in each pen *i* per cubic meter can be mathematically expressed as

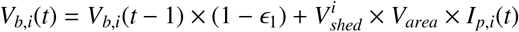

where *ϵ*_1_ denotes the decay rate of the viral particles. We used our data of pen-dimension (*V*_*area*_ defined in Table 1) to determine the level of viruses that will be present in each pen. The cumulative virus that can be transported to pen *i* from neighboring pens at each time step can be estimated as

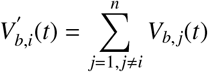

The following equation quantifies the likelihood of a pig in pen *i* getting Infected by this route:

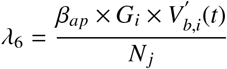

*G*_*i*_ is the cumulative virus that is transferred to pen *i* from all other pens. *β*_*ap*_ is the aerosol transmission rate as defined in section 2.3.3. *N*_*j*_ is the number of pigs in the neighboring pens. The contribution of room-level transmission contribution to disease transmission can be defined as

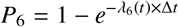

### 2.5 Human-mediated transmission at room level

In the ASFV transmission dynamics, Day-to-Day employees’ activities are related to the probability of disease dissemination [11, 58, 59]. Here, we modeled contaminated personnel entering barns and spreading the virus deterministically. The force of infection increases along with the number of Infected pigs in the room. For instance, the daily indirect transmission from humans (a.k.a. fomites, contaminated clothes, boots) to pigs on a farm is defined as,

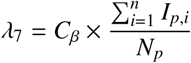

where *C*_*β*_ is the human transmission rate, *N*_*p*_ is the total number of pigs in the room, and *I*_*p*_ is the total number of Infected pigs in the room. Therefore, the probability that Susceptible pigs can get Exposed by this route is

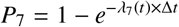

A schematic of the above transmission route is defined in Supplementary Material Figure S7.

Combining all within-pen and between-pen transmission routes, a Susceptible pig is likely to get Exposed as follows

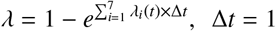

### 2.6 Moving pigs among pens

A regular practice in commercial swine production is sorting and moving animals among pens, mainly to improve production and group animals that are either sick or require additional treatment or feed [60]. Isolating sick animals significantly impacts pen-to-pen dissemination [16] while also allowing sick pigs to recover without competing with healthy animals. However, the transfer of pigs from one pen to another pen has also been associated with within-barn ASFV dissemination [9]. We generate a probability distribution function to simulate the daily pen-to-pen transferring of pigs to replicate swine production management.

We define the room within the barn as a directed graph *G* = (*V, E*), where pens are defined as a set of vertices *V* = *v*_1_, *v*_2_, …, *v*_*n*_, and *E* is the set of directed edges between vertices, where each edge *v*_*i*_, *v* _*j*_ represents a possible daily transfer of pigs from pen *i* to pen *j*. Each edge is associated with the transfer probability of *p*_*ij*_, where 0 ≤ *p*_*ij*_ < 1 is defined in Table 1 The process of transferring pigs is stochastic and modeled as follows:

1. A daily transfer random event consists of draws from a uniform distribution *U*(0, 1) for each edge *e*_*ij*_, a random number, *r*. If *r < p*_*ij*_, a transfer from pen *i* to pen *j* is executed.
2. If the transfer is executed, the number of pigs to be transferred is determined by a Poisson distribution with mean *λ* = 1, reflecting the expected number of pigs to transfer per event.
3. The selection of the source pen (from which pigs are transferred) and the target pen (to which pigs are transferred) is a random process.
4. The model restricts the daily number of transfer events to a maximum of three.
5. All pigs, regardless of their compartment, are equally likely to be chosen for transfer.
6. The model also ensures that pigs are not transferred back to the pen from which they were moved during the transfer process.

### 2.7. Outputs

Our model outputs include i) the infectious status of animals and pens over 175 Days; ii) the contribution of each transmission pathway to the overall transmission dynamic; iii) the viral load room distribution from fecal and aerosol transmission routes 175 Days; iv) pen status comprising fully Susceptible, Infected pens with no detection and at least one Detected pigs in a pen over 175 Days; v) the room distribution of Infected pens over time; and vi) pigs disease status distribution in a room over time. In addition, we developed a scenario-based analysis to evaluate variations in human-mediated ASFV transmission, mortality rates, and probability of pig transfers. For outputs i) ii), iii), and iv), we seed the infections randomly in one pen and examine the virus dissemination for 175 Days. For outputs v) and vi), we demonstrate the disease spread dynamics within a single room containing 32 pens. We ensured that each pen was the starting point of infection 100 times, given the 3200 repetitions of our model, to ensure that each pen was equally Exposed to the initial infection. For the scenario-based scenario, we consider two different rooms comprising 24 pens and 40 pens and vary the parameters.

## 3. Results

### 3.1. ASFV dissemination and contribution of transmission routes

On average, 50% of pigs became Exposed within 20 Days of ASFV introduction regardless of room sizes, while in 60 Days 95% were Exposed (Figure 3 A). The peak of newly Exposed pigs was at 17% (95% Confidence Interval (CI) = 16.9%17.1%) on Day 20, while for Clinical pigs was 3.93% (CI = 3.91%-3.95%) on Day 24, for Sub-Clinical pigs was 1.31% (CI = 1.29%-1.34%) on Day 24 and for Carrier pigs was 0.57% (CI = 0.55%-0.59%) on Day 24 (Figure 3 A). The cumulative prevalence indicated that 4.44% (CI = 4.34%-4.54%) of pigs exhibited Clinical signs within one week after the virus was introduced, 12.61% (CI = 12.5%-12.63%) after two weeks, and 36.83% (CI = 36.81%-36.85%) after one month. The cumulative prevalence of Sub-Clinical was 1.16% (CI = 1.03%-1.33%) after one week, 3.14% (CI = 2.94%-3.34%) after two weeks, and 8.77% (CI = 8.56%-8.98%) after one month. Carrier pigs showed the lowest cumulative prevalence, with 2.76% (CI = 2.68%-2.98%) after one month (Figure 3 C).

**Fig. 3:**
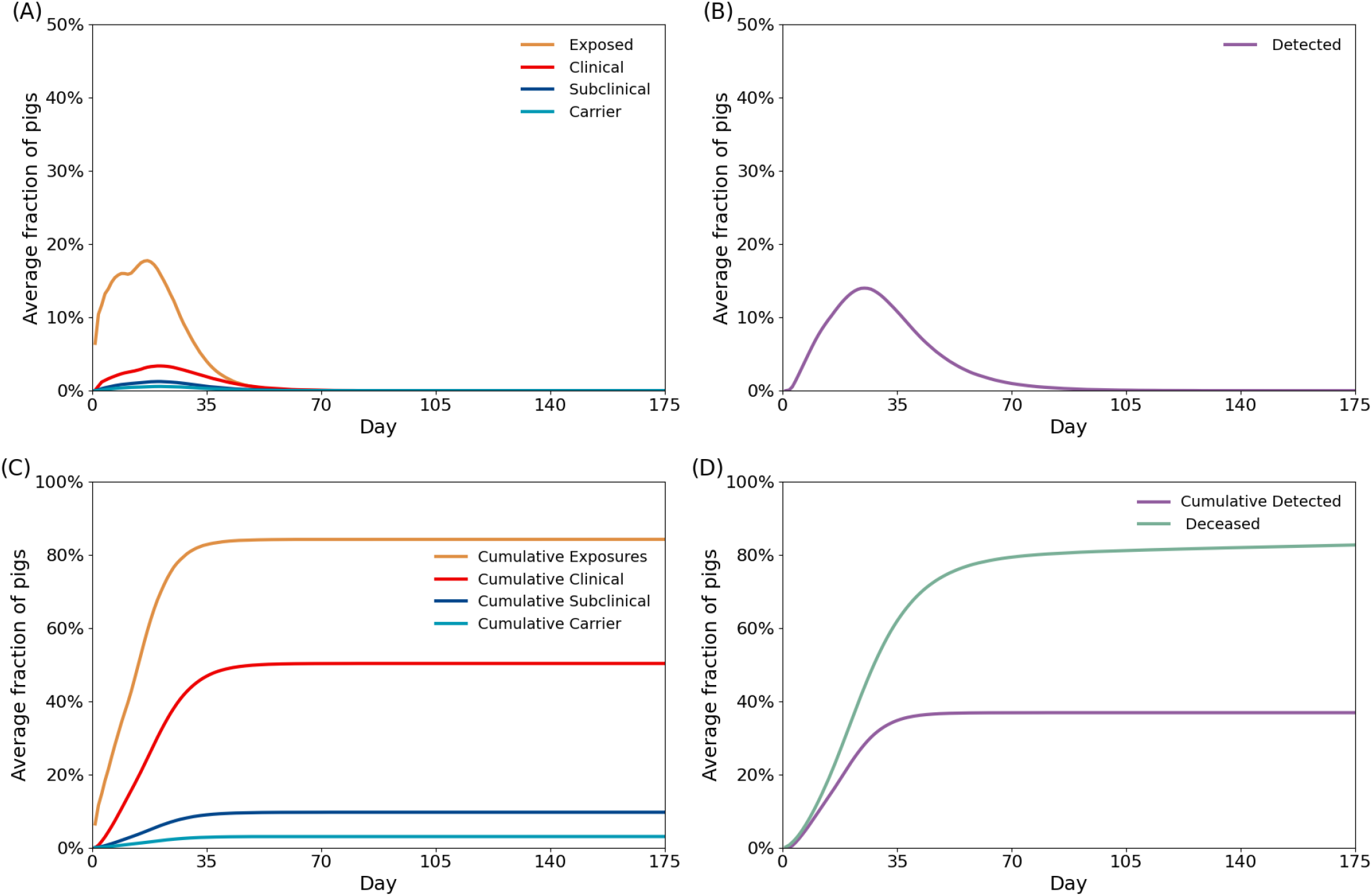
The upper panel illustrates the mean prevalence dynamics of ASFV spread over 175 Days across 7704 rooms. A) Mean prevalence of Exposed, Clinical, Sub-Clinical, and Carrier pigs, B) Mean Detected pigs over ASFV spread. The lower panel displays the cumulative disease prevalence of C) Exposed, Clinical, Sub-Clinical, and Carrier pigs, and D) Detected and deceased pigs. The y-axis represents the mean percentage of pigs. The Susceptible and Recovered populations are shown in Supplementary Figure S18.

The average time to detect the first Infected pig at room level was three Days (CI = 2-4 Days) (Figure 3 B). On average, 10% of pigs were Detected on Day 14, 25% were Detected on Day 27, and the maximum percentage of Detected pigs was 38% on Day 42 (Figure 3 B). The 10% cumulative mortality was assumed as a trigger value for active surveillance and was reached on Day 11 (CI = 10.66-10.87). We estimated that at the end of 175 Days, an average of 85.02% (CI = 84.86%-85.19%) pigs died, while 9.45% (CI = 9.38%-9.52%) of pigs recovered (Figure 3 D and Supplementary Material Figure S18).

The contribution of within-room transmission varied throughout the epidemic (Figure 4). In the first week, the between-pen fecal material was the main transmission route, contributing on average 45.97% (CI = 45.78%-46.12%) to the number of secondary infections, followed by direct contact of pigs between pens with 37.01% (CI = 36.94%-37.12%). Beyond week two of the simulation, the frequency of secondary infections due to direct contact with pigs between pens increased, while infections due to between-pen fecal material decreased (Figure 4). After 60 Days, the average contribution of direct contact of pigs between pens was 51.21% (CI = 51.11%-51.3%), and between-pen fecal material was 13.37% (CI = 13.27%-13.46%). Similar to the contact of pigs between pens, all other transmission routes gradually increased their transmission contribution over time, except for human-mediated transmission, which slightly decreased. The direct contact between-pen showed the highest contribution on Day 60 with 60.76% (CI = 60.68%-60.84%), followed by direct contact within-pen with 13.82% (CI = 13.74%-13.91%), fecal transmission within-pen with 10.9% (CI = 10.81%-10.98%), room-level airborne transmission with 6.37% (CI = 6.29%6.43%), and finally human-mediated transmission with 5.25% (CI = 5.19%-5.33%.) In our simulation, within-pen aerosol had minimal impact on disease spread, contributing to less than 1%.

**Fig. 4:**
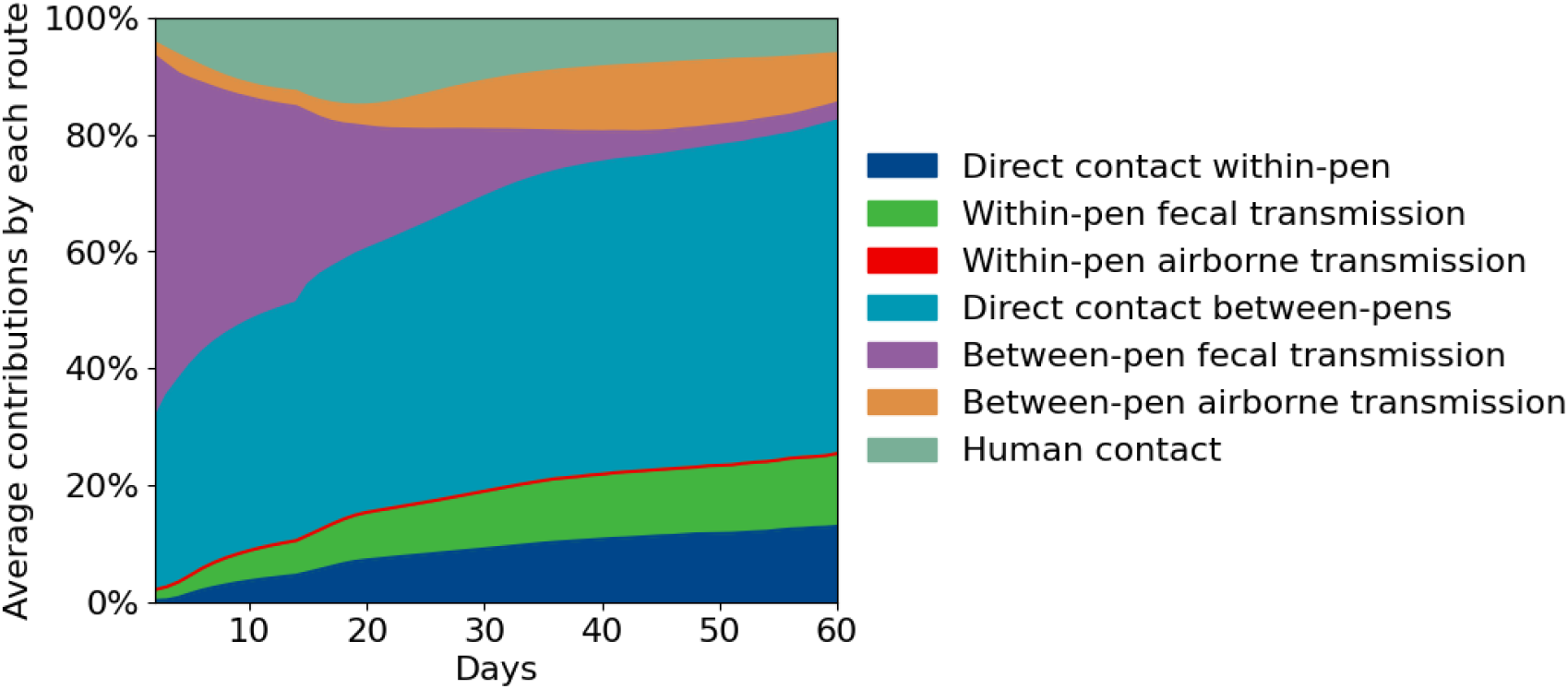
The average contribution of seven routes of within-room ASFV transmission over 60 Days (175 Days are not displayed as the number of Susceptible pigs across rooms approaches zero around 60 Days, resulting in almost no exposure events). The vertical axis represents the mean proportion of the contribution of each transmission route on a given Day. The data has been averaged at each time point and smoothed using a moving window approach.

### 3.2 ASFV overall dissemination at pen-level

Out of 7704 rooms to which ASFV was seeded, 58.06% (CI = 57.97%-58.14%) pens in each room were Exposed within one week of introduction, 82.59% (CI = 82.43%-82.76%) after two weeks, and 97.69% (CI = 97.58%-98.8%) after one month (Figure 5 A). We demonstrated that pens closer to the exhaust fan exhibited a higher frequency of infections in the first five to ten Days (Figure 6). We note similar results for the total dissemination within rooms; for instance, in a room with 28 pens, the distribution of Infected pens became homogeneous within 30 Days, with only pens 1 and 17 showing a lower frequency of infections. Pens with single neighbor pens (pen 1, pen 16, pen 17, and pen 32) tended to be less Infected in the early Days of infection. However, in 30 Days, pen 16 and pen 32, are highly Infected. On average, the first Detected pen was on Day 3 (CI = 2-4) (Figure 5 B), and the peak of detection was 81.04% (CI = 80.87%-81.12%) on Day 37 (Figure 5 B).

**Fig. 5:**
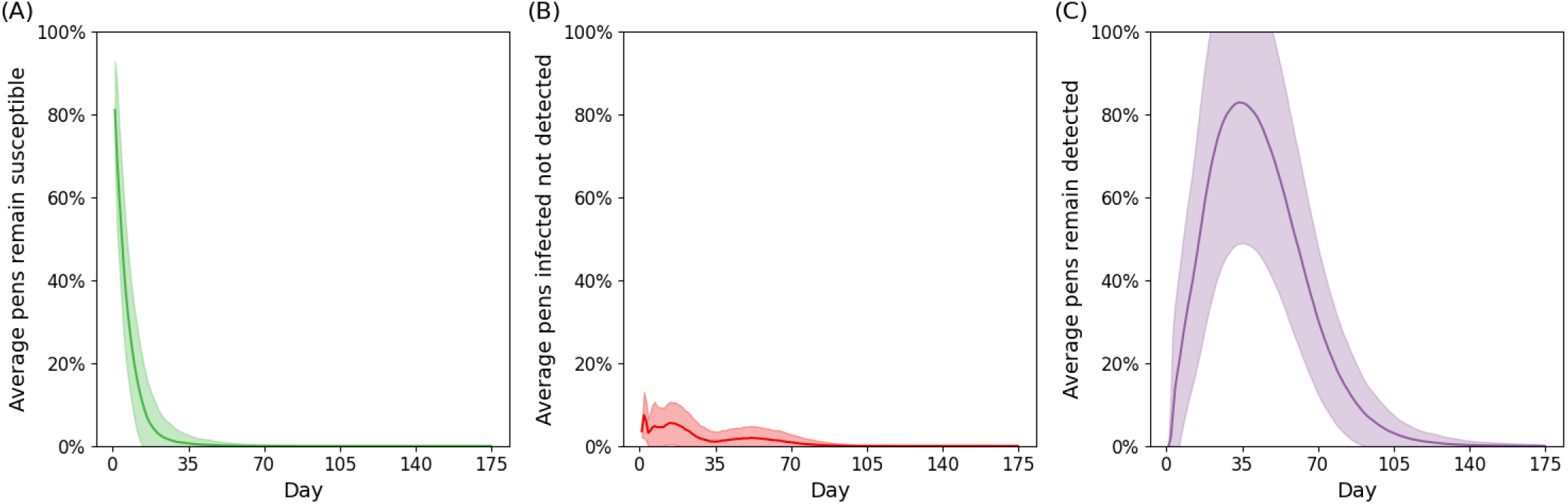
The daily average with a 95% confidence interval for the standard deviation of Susceptible, Infected, and Detected at pen-level. (A) Average percentage of pens with Susceptible pigs; (B) Average percentage of pens with at least one Infected pig; (C) Average percentage of pens with at least one Detected pig.

**Fig. 6:**
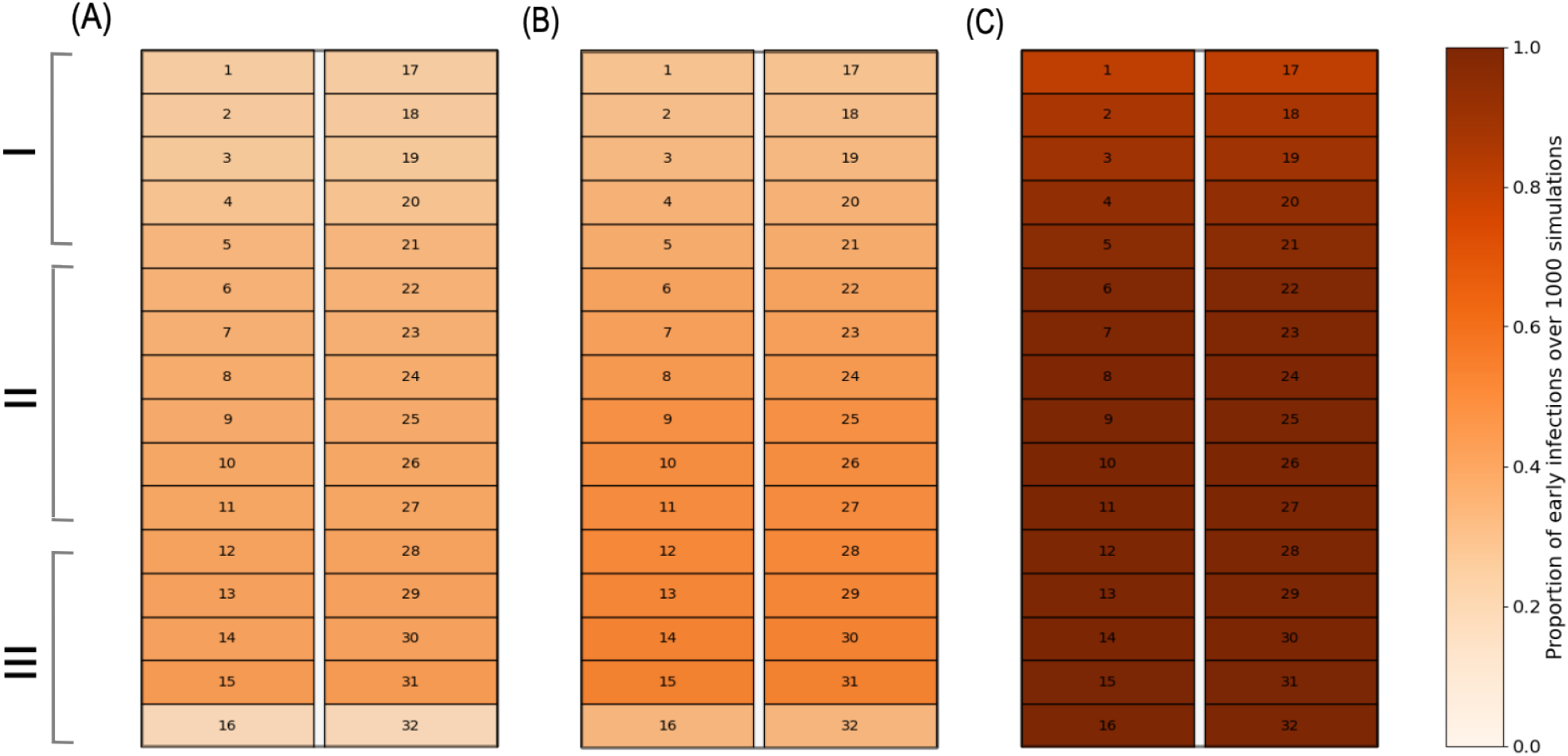
Number of times pens were identified as Infected at Day 5, 10, and 30 Days, respectively, for a room of 28 pens over 1000 simulations. The fans were placed near Pen 1 and Pen 15. The exhaust fans are located in Pen 14 and Pen 28. Description of the barn structure is defined in 1. We distribute the barn into three sections where I, II, and III.

We also evaluated the dynamic of Exposed, Clinical, SubClinical, and Carrier pigs across a room comprising 32 pens over three distinct time intervals: 5, 10, and 30 Days (Supplementary Figures S14-S18). The number of Exposed pigs in all pens decreases from Day 5 to Day 10 and continues to decline by Day 30 (Supplementary Material Figure S14). Most of the Exposed pigs were in room sections II and III during Days 5 and 10 but moved towards section I by Day 30 (Supplementary Material Figure S14). We also observed fewer Exposed pigs in sections I and II of the room on Day 30, as most had become Infected after the latency period. In contrast, the numbers of Clinical pigs (Supplementary Material Figure S15), Sub-Clinical pigs (Supplementary Material Figure S16), and Carrier pigs (Supplementary Material Figure S17) increased linearly from Day 5 to Day 30. By Day 30, all pens had a higher number of Infected pigs, including Clinical, Sub-Clinical, and Carrier pigs. The above trend could be attributed to the incubation period of Exposed pigs and their progression to Clinical, SubClinical and Carrier pigs as they complete the latency period. The sections that were initially Exposed on Day 5 are the ones that now contain Clinical pigs.

### 3.3. Viral load dynamics of fecal and aerosol transmission

Fecal material viral load was 2 ×10^6^ time higher than airborne viral load (Supplementary Material Figures S13 (A) and S13 (B)) across all rooms. The average viral load from fecal material and aerosol peaked on Day 28. Following the peak, the viral load stemming from fecal matter and aerosols steadily diminished, gradually approaching zero towards Day 175, which could be attributed to the decline in Infected pigs (Figure 3). The mean viral loads generated from fecal (*Q*_*a*_) accumulation at pen level, is 5 ×10^7^ faster than aerosol (*V*_*b*_) (Figure 7). The viral loads on pens from both fecal and aerosol sources showed a similar distribution at the room level in the first 30 days after the virus was introduced. Higher viral loads were found in pens near the far end of the room (Figure **??**, pens in Section III for a room with 24 pens, and Supplementary Material Figures S12 for a room with 40 pens), close to the exhaust fans. Conversely, pens with lower viral loads were located near the entry of the room (Figure **??**, pens in Section I for a room with 24 pens, and Supplementary Material Figures S12 for a room with 40 pens).

**Fig. 7:**
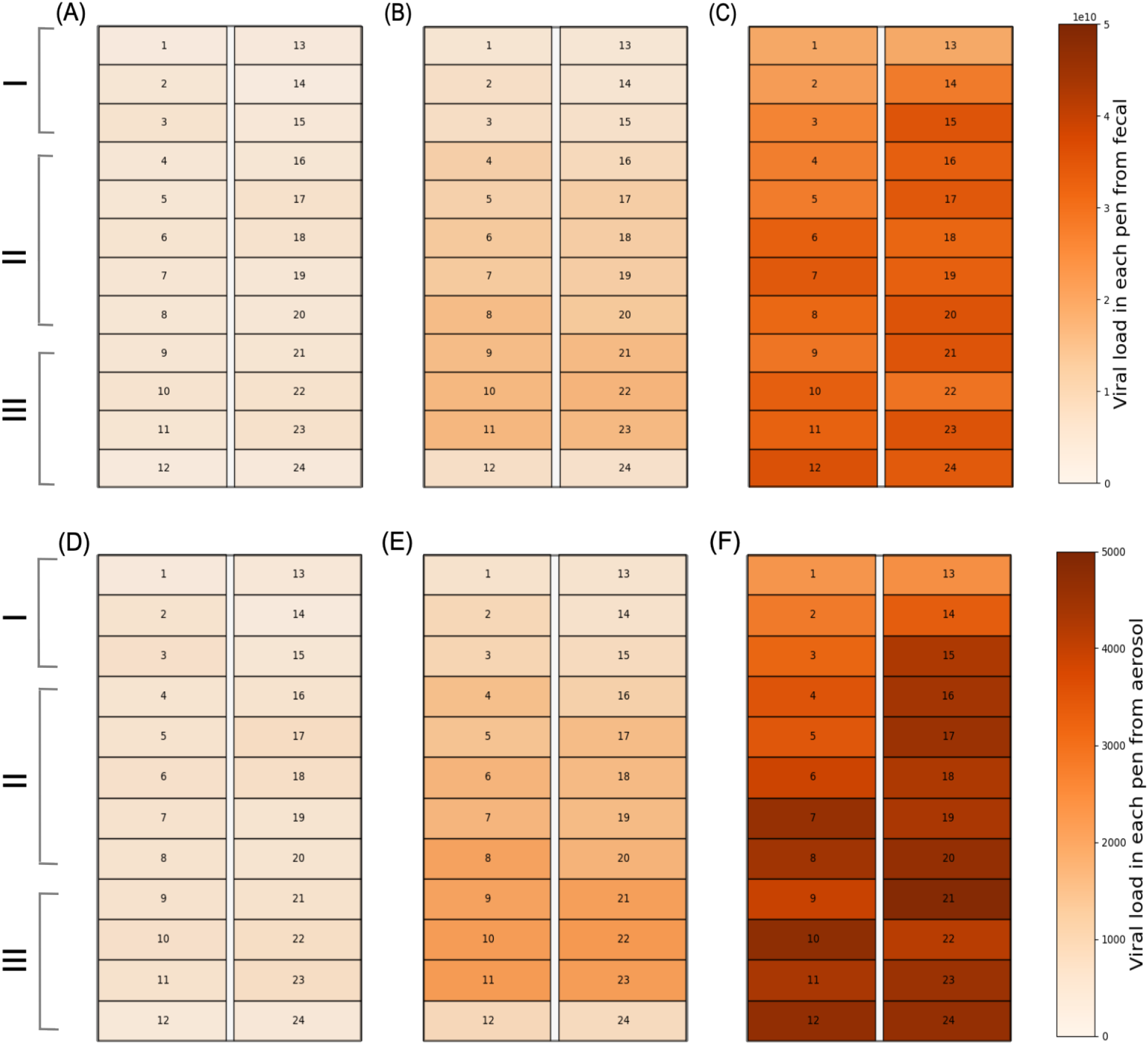
Average viral loads generated through fecal transmission (top panel) and aerosol transmission (bottom panel) across 24 pens over periods of (A, D) 5, (B, E) 10, and (C, F) 30 Days.

### 3.4. Scenario-based and sensitivity analyses

#### 3.4.1. Pen-to-pen pigs transferring

We examined the effects of varying the frequency of transferring pigs between pens on ASFV spread. We used daily pig transfer probabilities at 5%, 25%, and 50% for a room containing 24 pens and 40 pens. Supplementary Material Figures S8 and S9 demonstrate that altering the probabilities of pig transfers does not impact significantly ASFV dissemination dynamics. The contribution of each transmission pathway remains similar across the three different scenarios.

#### 3.4.2. Human-mediated transmission

We investigated the impact of the human-mediated transmission parameter by varying it fivefold and tenfold in a room comprising 24 pens, simulating increased human visits to each pen. As the human-mediated transmission rate increased, the peak of the outbreak was observed earlier. For instance, in Figure 8 A), which represents our base model, the peak for Exposed pigs occurred on Day 17 and for Clinical, Sub-Clinical, and Carrier pigs on Day 19. When the human-transmission rate was increased fivefold, the peaks for Exposed pigs shifted to Day 14 and Clinical, Sub-Clinical, and Carrier pigs shifted to Day 17, respectively, as shown in Figure 8 B). With a tenfold increase, these peaks shifted further to Day 11 for Exposed and Day 14 for Clinical, Sub-Clinical, and Carrier pigs, respectively (Figure 8 C).

**Fig. 8:**
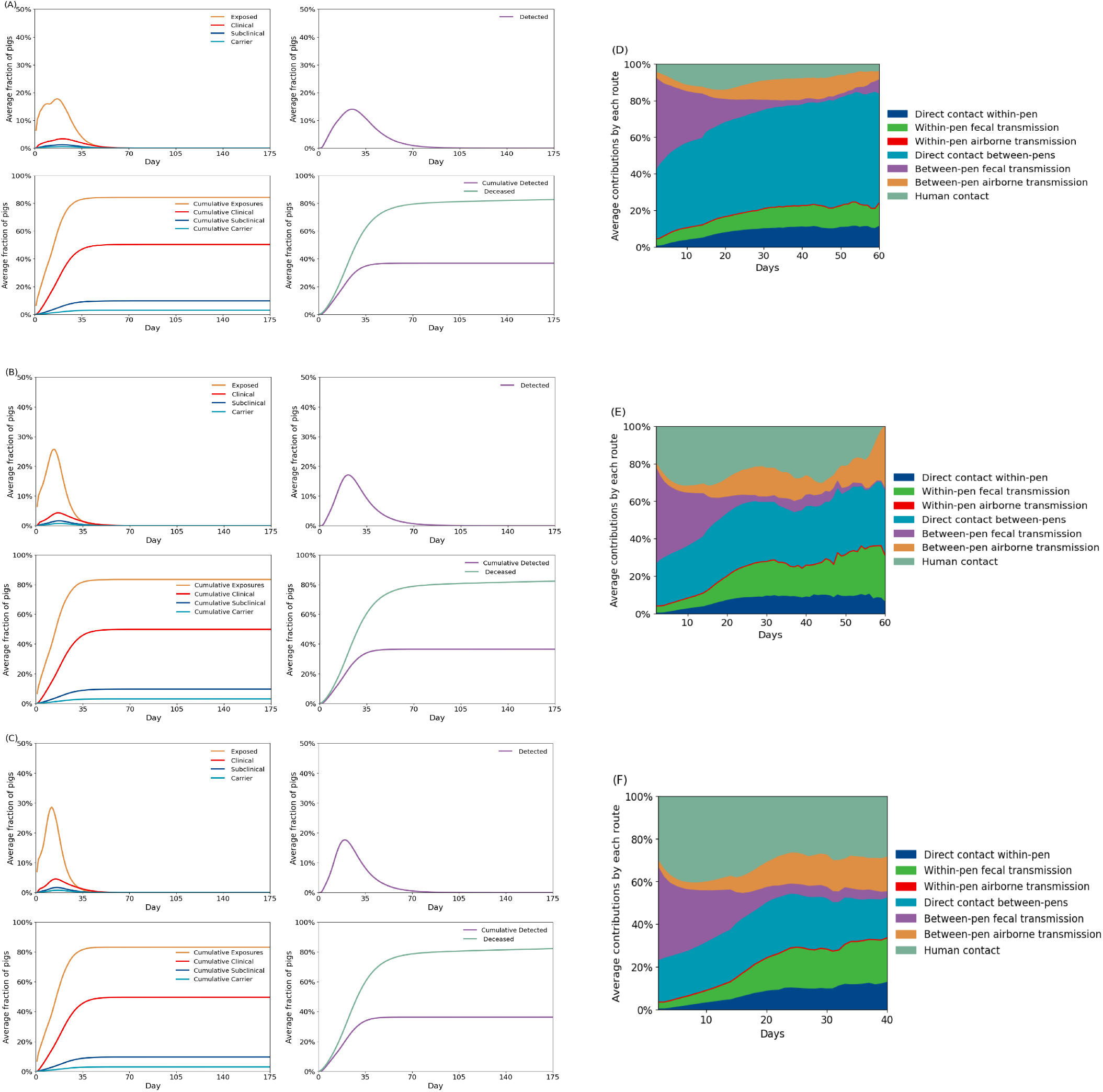
We investigate the spread of ASFV and the contribution of individual routes in a room consisting of 24 pens under different human-mediated transmission scenarios: A) and D) for the base model with *C*_*β*_=0.01, B) and E) for *C*_*β*_ =0.05, and C) and F) for *C*_*β*_ =0.1.

As the contribution of the human-mediated transmission route increased fivefold and tenfold, the average contribution of within-pen routes (direct nose-to-nose contact and fecal transmission within pens) also increased, as this route enhances disease spread within a pen (Figures 8 E and 8 F). Furthermore, in the scenario with the highest human-mediated transmission, all Susceptible pigs were Exposed within 40 Days (Figure 8 F), compared to 64 Days for the medium transmission rate (Figure 8 E) and 90 Days in the base model (Figure 8 D). We also observed similar dynamics in scenarios with rooms with 40 pens (Supplementary Figure S10).

#### 3.4.3. Mortality rates

We analyzed ASFV dynamics under varying mortality rates by both decreasing and increasing the mortality rate five times for pigs in different compartments in a room comprising 24 pens. Higher mortality rates led to a reduction in the overall number of Infected pigs, thus decreasing the cumulative numbers of Exposed, Clinical, Sub-Clinical, and Carrier pigs. For the base mortality rate as shown in 1, the cumulative Exposed pigs were 82.75% (CI = 81.69%-83.63%), Clinical pigs were 49.4% (CI = 48.54%-50.34%), Sub-Clinical pigs were 9.58% (CI = 8.65%-10.48%), and Carrier pigs were 3% (CI = 2.92%-3.09%), Figure 9 A). When we reduced the mortality rate by five, the cumulative Exposed pigs were 89.58% (CI = 87.91%-91.24%), Clinical pigs were 70.22% (CI = 68.91%-71.52%), Sub-Clinical pigs were 13.61% (CI = 13.35%-13.87%), and Carrier pigs were 4.11% (CI = 4.02%-4.19%), Figure 9 B). Conversely, when we increased the mortality rate by five times compared to the base model, the cumulative Exposed pigs were 28.2% (CI = 26%-30.4%), Clinical pigs were 7.79% (CI = 7.18%-8.41%), Sub-Clinical pigs were 1.47% (CI = 1.35%1.59%), and Carrier pigs were 0.48% (CI = 0.48%-0.51%) (Figure 9 A).

**Fig. 9:**
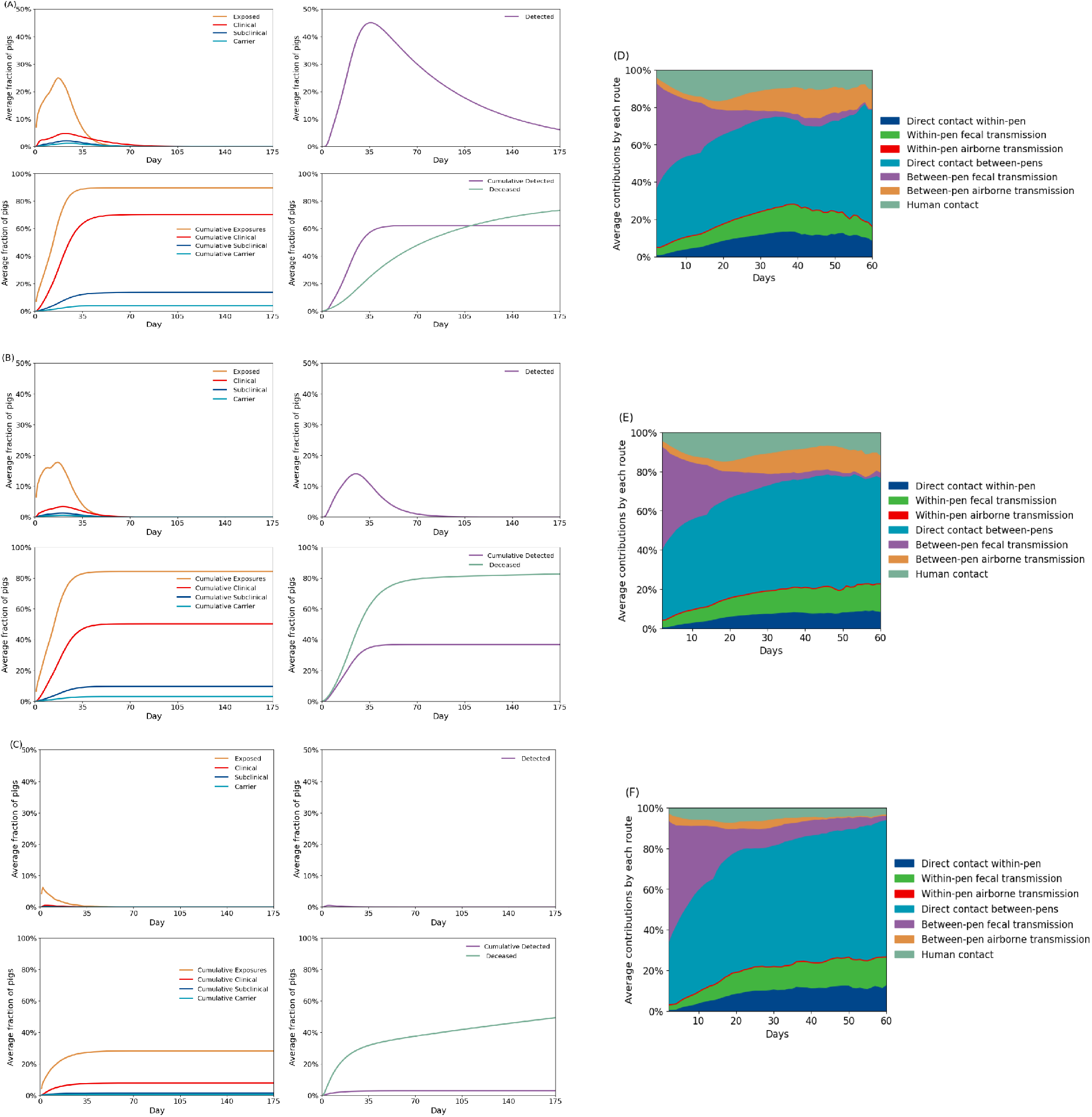
We investigate the spread of ASFV and the contribution of individual routes in a room consisting of 24 pens under different mortality rates: A) and D) for the low mortality rate where the mortality rate is reduced by five times, B) and E) for base mortality rate and C) and F) for the high mortality rate where the mortality rate is increased by five times

Additionally, the number of Detected pigs reduced as the mortality rate increased. For the base model, the cumulative Detected pigs were 36.18% (CI = 35.23%-37.13%) (Figure 9 A), whereas for lower mortality rates, the cumulative Detected pigs were 62.24% (CI = 61.08%-63.4%) (Figure 9 B), and for higher mortality rates, the cumulative Detected pigs were 2.82% (CI = 2.6%-3.04%) (Figure 9 C). Interestingly, changes in the mortality rate had only a marginal impact on the contribution of different routes to disease spread (Figures 9 D-F).

## 4. Discussion

We have developed a stochastic multilevel population model to enhance our understanding and make projections about the spread of ASFV within commercial swine barns. This model allows us to analyze the dissemination of the virus both within the overall population and at the pen level, tracking how pens and pigs become Infected. In the first week of simulations, 21.8% of the total animals and 58.06% of the total pens had Infected animals on average, underscoring the rapid within-room dissemination of ASFV. After one month, nearly all animals and pens were Infected, demonstrating the virus’s capacity to sustain transmission over a prolonged period and affect nearly the entire room population. For instance, within the first two weeks, 24.96% of animals were Exposed while 27.19% of pens remained Susceptible. The model also indicates that fecal transmission between pens was the primary route of infection in the first week, but this was later surpassed by direct contact between pigs starting from the second week. Notably, most Infected pens were initially located in the middle of the rooms at five and ten Days, with direct contact and fecal transmission being the main contributors to disease spread. The proximity to multiple neighboring pens increases the likelihood of disease spread through direct contact and fecal transmission [23], as these pens are more frequently Exposed to Infected pigs and contaminated materials from adjacent pens (Figure 4). As time progressed, proportions of pens near the exhaust fans also became Infected. This can be attributed to increased exposure of pigs to contaminated air during this period around the exhaust fans, leading to a rise in aerosol transmission between pens, which depends on the total number of infected pigs within the room(s) (Figures 6 and 7).

The average time to ASFV room-level detection was on Day three, with an associated average pig mortality rate of 0.64% (CI = 0.6%-0.68%). This early detection is attributed to passive surveillance, which was 1% in our current model, in identifying initial infections. As the disease spread, the mortality rate increased, prompting an intensification of active surveillance. By Day 14, approximately 10% of pigs were Detected, and the detection rate peaked at 38% on Day 42 (Figure 3 D), primarily driven by active surveillance. This active surveillance method became more effective in identifying and responding to the outbreak as mortality escalated or exceeded 10% of the total population daily. These findings highlight the necessity of combining passive and active surveillance to manage the disease effectively as it progresses. Given the challenges of early ASFV detection [3, 61, 62] and catastrophic propagation consequences within swine farms [63, 64], our findings highlight ASFV transmission dynamics at room and pen levels and provide valuable insights to strengthen surveillance strategies at a wide range of barn layouts.

Over 60 Days of within-room spread, we observe that between-pen transmission pathways contributed more than 71% on average towards dissemination, while the within-pen transmission pathways contributed about 21% on average. The significance of between-pen transmission can be attributed to the homogeneous pen configuration as pens have similar dimensions and are closely spaced, thereby enhancing pen-to-pen disease transmission, as shown elsewhere [65]. Furthermore, by introducing a single infected pig randomly into one pen, we ensure that for direct nose-to-nose contact and fecal transmission pathways (both within-pens and between-pens) to be more pronounced, the pigs need to come into contact with neighboring Infected pens. The easy spread of the virus between pens highlights the critical need for targeted pen-level measures. For instance, removing Infected pigs from pens is crucial to reducing the probability of direct contact transmission. Enhanced surveillance strategies, such as regular monitoring and testing of pigs in pens located in rooms’ sections II initially (Figure 6), could be utilized for targeted surveillance. Environmental sampling, including testing surfaces and air samples, could also help identify viral presence and thereby implement strategic interventions [15, 20, 66, 67].

Within-room environmental contamination via fecal and airborne routes accounted for more than half of transmissions in the first week on average. This proportion decreased over time, representing 31.23% of transmissions by Day 60. These results highlight the importance of environmental contamination at the initial stage of ASFV propagation [30, 68], and to continue infecting pigs over time [30, 69]. Strategies to mitigate fecal contamination, such as thorough cleaning and disinfection, along with effective ventilation or air filtration systems, could significantly curb ASFV dissemination [70, 71]. Transmission by direct contact of pigs within and between pens constantly increased after the first week, reaching more than half at Day 60. Currently, removing Infected pigs from pens is the only strategy that reduces the probability of transmission by direct contact [66, 67]. However, this strategy is strongly linked to the capacity for early detection of Infected pigs to reduce the time that naive pigs are Exposed [64].

The human-mediated introduction of diseases into commercial swine farms is recognized as a significant pathway for disease entry [72]. However, its role in pen-to-pen dissemination remains to be examined. Contrary to the other transmission routes, human-mediated transmission showed more variable patterns over time, with a peak of transmission on Day Although this transmission route exhibited a lower contribution compared with environmental contamination and direct contact (Figure 4), the human transmission plays a crucial role in spreading the virus among rooms, barns, and farms [59, 64]. Our findings demonstrate that increasing the transmission rate parameter associated with human-mediated introduction can lead to faster within-room dissemination of ASFV. This is because humans travel between pens, potentially spreading diseases to unaffected pens. Biosecurity strategies have been shown to reduce the probability of introduction; measures such as restricting access to essential personnel, establishing clear boundaries between clean and dirty areas, and requiring decontamination procedures, use of a perimeter fence, and designated parking areas could also minimize disease transmission. Additionally, hygiene practices, such as changing clothes and boots and using disinfectants, are crucial in preventing the spread of pathogens [67, 73].

Our findings indicate that transferring pigs between pens did not impact the spread intensity of within the barn spread. This is primarily due to the virus’s high baseline transmission rate, allowing it to spread rapidly and efficiently through various other pathways, regardless of rearranging pigs between pens. Furthermore, due to indirect transmission from fecal and aerosol contamination, the virus is able to spread widely early on the outbreak, so moving pigs between pens does not necessarily introduce them to new sources of infection that they have not already encountered. In our analysis, the objective of moving pigs is for higher productivity and control aggressive and sexual behavior for better animal welfare as seen in commercial farm practices, and therefore pigs are moved regardless of their disease status [74, 75]. However, if the indirect transmission from fecal and aerosol is not widespread, moving Infected pigs between pens could be a crucial route in transferring infections to unaffected pens, thereby increasing disease spread. While the literature remains controversial on the roles of Clinically Infected pigs and Sub-Clinically Infected pigs in the dissemination of ASFV, our results showed that Clinical cases significantly contribute to the spread of ASFV due to higher virus shedding rates and observable symptoms, thus providing insight into this complex dynamic within commercial swine farms in the U.S. [3, 76, 77], conversely, studies by [23, 24] highlight the under-recognized yet critical role of Sub-Clinical infections. Sub-Clinical pigs can act as Carriers, facilitating the unnoticed spread of the virus [66, 78]. Our results demonstrate that when the mortality threshold crosses 10%, thereby triggering active surveillance, clinical animals would represent 80% of the room population, while sub-clinical animals and carriers represent 15% and 5%, respectively.

Furthermore, ASFV is known for its high mortality rate when first introduced to naive populations. Our results indicate that the rapid increase in mortality among infected pigs limits the spread of the virus, as it reduces the opportunities for further transmission [43], Figure 9 (C). Our results clearly demonstrate a correlation between high mortality rates and decreased levels of exposure; as the mortality rate increases, the number of pigs that remain Infected long enough to spread the virus decreases. This leads to fewer Exposed cases over time since the low survivability of Infected pigs prevents the establishment of a larger pool of Infected pigs. In essence, while ASFV’s high mortality rate makes it a devastating disease, it also curtails the virus’s ability to spread extensively within barns.

## 5. Limitation and further remarks

At the time of writing this manuscript, the U.S. is ASFV-free [79], and current control strategies include stamping out the entire population of any Infected farm [80]. One limitation of this study was that it did not consider control actions. Indeed, this work aimed to describe the dynamics of ASFV at a wide range of barn sizes and layouts and quantify transmission routes. In the future, model interaction should consider the implementation of control actions at multiple levels, barn, room, and pen levels, for instance, simulating the effectiveness of barn or pen-level test and removal [81]. Additionally, the model should be expanded to consider the dissemination between rooms as a barn space is considered before the spread of diseases to be in the same air space, allowing dissemination via several modes [82]. Also, we did not account for ASFV stability in the environment[51, 68, 83]. Future models should include ASFV environmental stability to simulate residual contamination better and assess repopulation plans’ effectiveness [84]. Along these lines, because the regular finisher production cycle in North America is approximately 112 Days, we did not consider the introduction of new animals from new movements to our premises, as most farms are all-in and all-out sites, meaning that once pigs are harvested, all animals are removed and the farm remains empty for at least one Day before placing new animals [85, 86].

Through (RABapp^™^)[44], each farm has map that includes the dimensions of barns [44], which were used to derive the number of pens per room, based on literature [45] and our own field observations of pen dimensions. Given the heterogeneity among commercial swine barns’ internal layouts, the room and pen design distribution in our model may not precisely represent the actual barn sizes. However, our model validation indicates that it provides a good approximation to represent commercial barns in the U.S. In addition, pigs are frequently moved between pens and rooms due to group management (e.g., keeping same-size pigs in the same pens) or for disease management purposes [74, 75, 87]. Future iterations of our model will include the movement of pigs between rooms since the transfer of pigs between different rooms could significantly contribute to the speed of dissemination.

The absence of specific data regarding the frequency of personnel visits per pen presents a challenge in accurately calibrating transmission parameters related to human activity. Alternatively, we have approximated the human transmission parameter by using a fraction of the parameter assigned to direct contact transmission. Nevertheless, we conducted a sensitivity analysis of this parameter to assess its impact on diseas spread and the relative contribution of different pathways. Future research should aim to monitor and record the daily number of staff visits to each pen and estimate the rate at which it is spreading the disease, thereby generating more precise values for this parameter.

Despite the limitations of our model and data, our study simulates the spread of ASFV utilizing barn and room layouts of 1958 premises and 7704 rooms, thus approximating the expected dissemination of ASFV within U.S. commercial swine farms. This extensive dataset allowed us to create a transmission model that integrates the structure and production conditions of swine farms, providing a significant advantage in simulating the dynamics of infectious diseases under real conditions. As a result, our study offers more robust and detailed insights into the effects of barn-level surveillance. In addition, our within-room dynamic system considers eight transmission pathways, providing swine producers and animal health officials with critical insights into the dynamics of ASFV spread when introduced in commercial swine production. Furthermore, we have examined pen distribution and how sorting pigs between pens can significantly impact the spread of disease, thereby highlighting patterns that could be prioritized once ASFV is Detected in commercial swine premises in the U.S.

## 6. Conclusion

In this study, we analyzed the spread of disease within barns subdivided into rooms and further into pens by examining interactions between pigs within and between pens, as well as the overall ASFV dissemination at both the pen and room level. We identified direct contact and fecal transmission from adjacent pens as the primary routes for ASFV spread, emphasizing the importance of close proximity in pens on commercial farms for facilitating disease transmission. Our findings indicate that a heterogeneous pen structure significantly limits ASFV spread when transmission is confined to within-pen interactions, contributing an average of 20.1% to disease spread. Although this contribution is substantial, it is not the primary pathway for within-room transmission. In contrast, interactions between pens, driven by direct contact, fecal contamination, and aerosols, are crucial, contributing up to 71.4% on average to ASFV spread. We identified that ASFV could be Detected within three Days, with peak detection achieved when the mortality rates peaked at Day 42. Furthermore, we identified that the distribution of Infected pigs within rooms was not homogeneous; pens near the exhaust fan, showed a higher prevalence of Infected pigs. Moreover, the employment of a frequencydependent model in this study shows how the presence and number of Infected pigs critically influence disease dynamics, affecting both the peak and duration of disease prevalence. Overall, this research not only deepens our understanding of ASFV transmission dynamics at the pen and room level but also explains the critical roles of commercial swine barn practices, pen structure, and disease dissemination by various pathways of ASFV spread. Understanding these pathways should be helpful in prioritizing intervention strategies to control the spread.

## Footnotes

1 see https://www.securepork.org/

