## Supplementary material for "Modeling the transmission dynamics of African swine fever virus within commercial swine barns: Quantifying the contribution of multiple transmission pathways": SS

### Supplementary Material:Multiple routes for direct or indirect transmission of African swine fever virus into a barn

#### 1 Description of the methodology used to distribute pens within the 7704 rooms in our database

The LOS feature polygon boundary represents a physical barrier, typically the walls of barns. The LOS polygons were trimmed to create polygons that represent individual buildings. Polygons that were too small or irregular to represent a barn housing animals likely were removed. The remaining polygons were rectangularized in a way that minimizes distortion of the area and aspect ratio of the original polygon. These rectangularized polygons were then used to create polygons representing the rooms of all barns in our database. If the length of the shorter side of the barn rectangle was less or equal to 20 meters, then one room was created (a copy of the barn rectangle). If the length of the shorter side of the barn rectangle was greater than 20 meters, the barn rectangle was divided into two rooms by bisecting the shorter side of the rectangle. To create pens in the rooms, each room was first bisected with a 1-meter-wide walkway from the short side to the short side. The remaining area was divided into an even number of pens that were as close as possible to 25 square meters in area and symmetrically arranged on either side of the walkway. Fans were placed along the polygon boundaries representing the walls of the barns. A 1.2-meter-long fan was placed on each of the longer walls of the barns, at the corner with the shorter wall. 1.45-meter-long fans were placed on the opposite short wall; for each room, four fans were placed along this wall, with a fan at each corner of the room boundary and the other two fans placed such that there was equal spacing between the fans. With standardized pen sizes prevalent in the industry (averaging 19.6 ft $\times$ 9.8 ft, accommodating on an average 15 to 35 pigs per pen, we used the pen area as input to design and distribute pens within each room.

##### 1.1 Description of the methodology used to distribute pigs per pen

To determine the number of pigs per pen, we divide the total farm capacity by the product of the number of barns, the number of rooms per barn, and the number of pens per room. This can be expressed mathematically as:

$$\text{Pigs per pen} = \frac{\text{Farm capacity}}{\text{Number of barns} \times \text{Number of rooms per barn} \times \text{Number of pens per room}} \tag{1}$$

For instance, Farm 00F3M7B has a capacity of 8,640 pigs, with 8 barns, each containing 1 room and 34 pens. Using the formula, the average number of pigs per pen is approximately 32 (31.74).

### 2 Schematic of the seven transmission routes

This section provides a detailed schematic of the seven primary transmission routes that facilitate the spread of ASFV within a room. Our within-barn transmission model considers the spread of ASFV at three levels: 1) within pens, 2) between pens, and 3) at room level. In the following section, we illustrate the within-pen transmission of ASFV spread.

#### 2.1 Within-pen transmission

In this sub-section, we illustrate the following three pathways that define the spread of ASFV within a pen.

##### 2.1.1 Within pen nose-to-nose contact

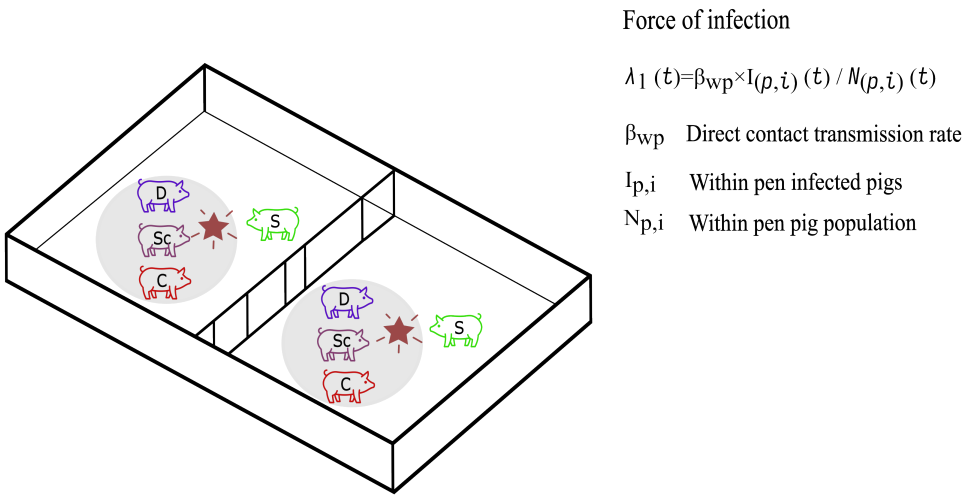


Figure 1: Schematic depicting direct nose-to-nose contact between pigs within a pen, demonstrating the pathway of direct disease transmission between infected and susceptible pigs sharing the same pen. $\beta_{wp}$ represents the direct contact transmission rate. $N_{p,i}(t)$ and $I_{p,i}(t)$ represent the total number of pigs and the number of infected pigs in pen $i$ at time $t$, respectively.

##### 2.1.2 Within pen oro-fecal transmission

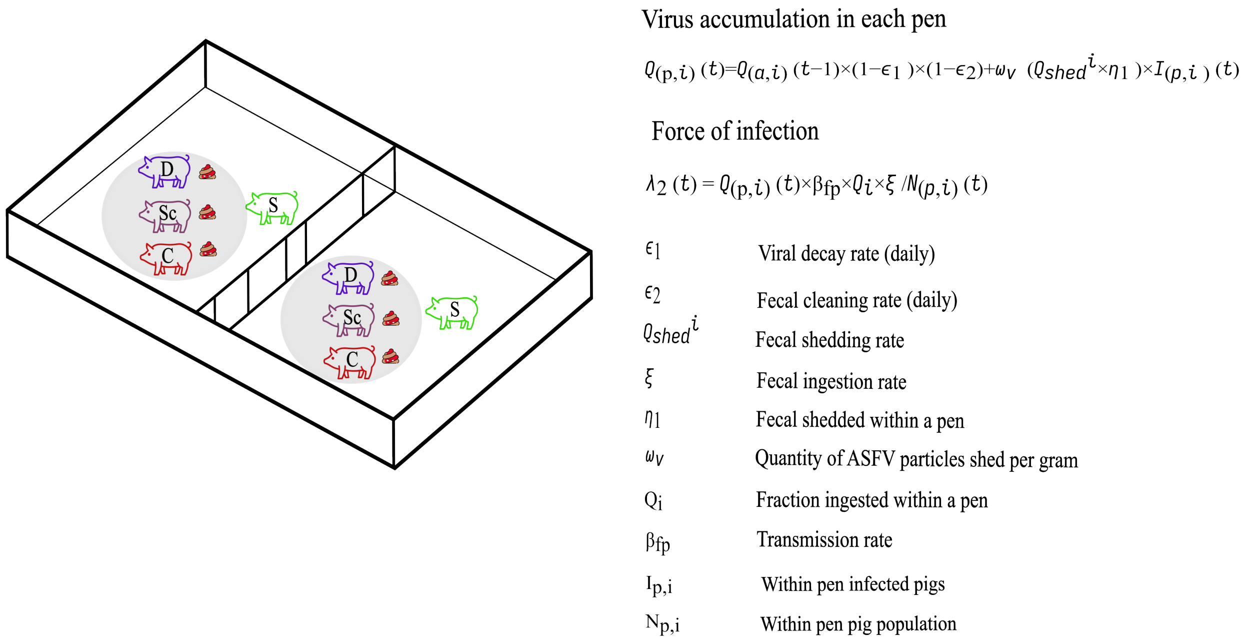


Figure 2: The schematic illustrates oro-fecal transmission within a pen, which is influenced by the daily accumulation of the virus due to the shedding of fecal matter by infected pig(s). The parameter $\omega_v$ represents the average amount of virus shed by an infectious pig per gram of feces each day and is defined as a uniform distribution spanning the minimum to maximum viral loads. The cumulative viral load depends on the average fecal load generated by an infectious pig daily, denoted as $Q^i_{shed}$. We also consider fecal shedding within and into adjacent pens, captured by $0 < \eta_1, \eta_2 < 1$, where $\eta_1$ and $\eta_2$ indicate the fractions of fecal matter shed within the pen and into adjacent pens, respectively. Additionally, we incorporate $\epsilon_1$ and $\epsilon_2$ to account for the daily removal of fecal matter through slatted floors and the environmental decay rate of ASFV, respectively. $Q_i$ is the daily amount of feces ingested by an animal. We consider that pigs ingest a fraction of the fecal denoted as $0 < \xi_1 < 1$ within the pen and the remaining $0 < \xi_2 < 1$ from its neighboring pens. $N_{p,i}(t)$ and $I_{p,i}(t)$ represent the total number of pigs and the number of infected pigs in pen $i$ at time $t$, respectively.

##### 2.1.3 Within pen air-flow transmission

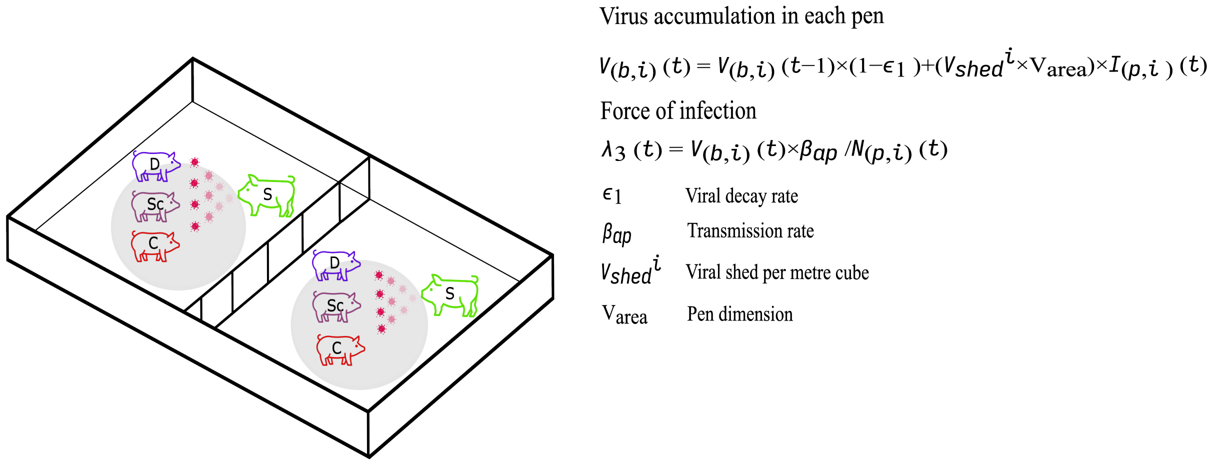


Figure 3: This schematic illustrates how aerosol transmission occurs within a pen. Infection by this route is driven by the daily accumulation of the virus in each pen, stemming from the virus shed by infected pigs wherein the virus is influenced by the daily decay rate of viral particles. We consider that the virus decays at a fixed rate ($\epsilon_1$) within a pen. The shedding rate of aerosol virus per cubic meter of the air by an infectious pig, $V^i_{shed}$, is characterized by a uniform distribution that encompasses the range from the minimum to the maximum viral load shed. We used our observed pen dimensions ($V_{area}$) to estimate the total viral load present in the air within each pen. $\beta_{ap}$ represents the direct contact transmission rate. $N_{p,i}(t)$ and $I_{p,i}(t)$ represent the total number of pigs and the number of infected pigs in pen $i$ at time $t$, respectively.

#### 2.2 Between-pen transmission

In this sub-section, we illustrate the following three pathways that define the spread of ASFV between adjacent pen.

##### 2.2.1 Nose-to-nose pig contact between adjacent pen(s) transmission

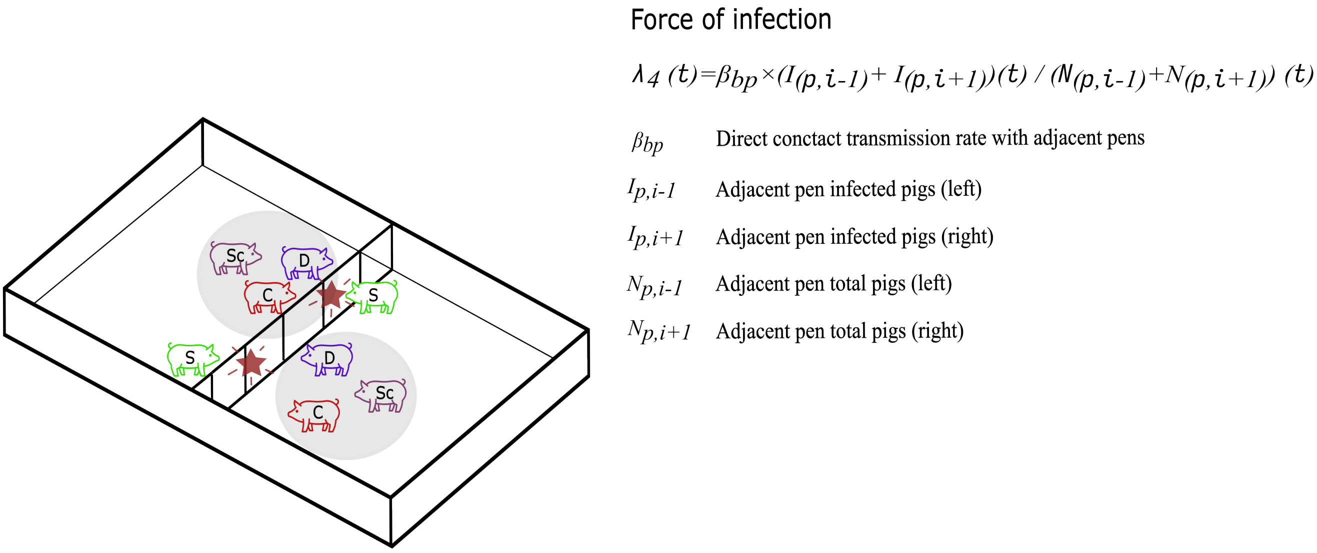


Figure 4: The schematic illustrates direct nose-to-nose contact between pigs in adjacent pens. It takes into account the number of infected pigs in neighboring pens to estimate the force of infection. Here, $\beta_{bp}$ represents the virus transmission rate, while $N_{p,i-1}(t)$, $N_{p,i+1}(t)$, $I_{p,i-1}(t)$, and $I_{p,i+1}(t)$ denote the total number of pigs and the number of infected pigs in pen $i-1$ and pen $i+1$ at time $t$, for the left and right neighbors, respectively.

##### 2.2.2 Between pen oro-fecal transmission

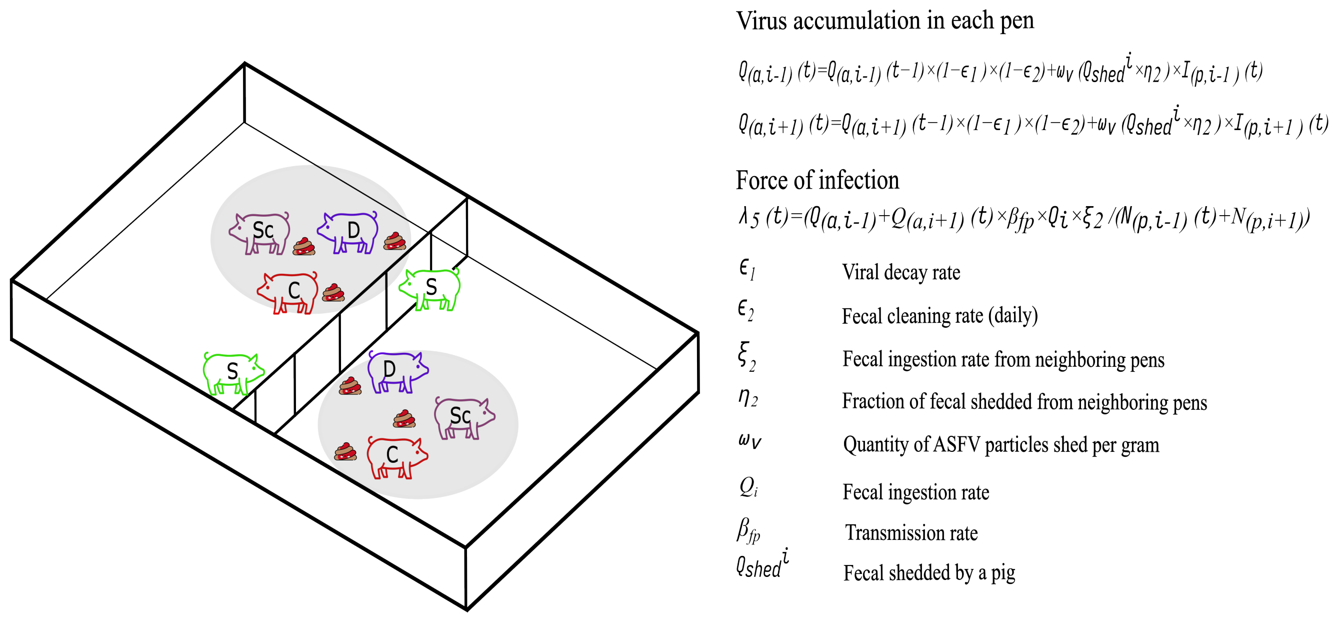


Figure 5: Schematic depicting the route of fecal transmission between pens. Here $\omega_v$ represents the average amount of virus shed by an infectious pig per gram of feces each day and is defined as a uniform distribution spanning the minimum to maximum viral loads. The cumulative viral load depends on the average fecal load generated by an infectious pig daily, denoted as $Q^i_{shed}$. We also consider fecal shedding within and into adjacent pens, captured by $0 < \eta_1, \eta_2 < 1$, where $\eta_1$ and $\eta_2$ indicate the fractions of fecal matter shed within the pen and into adjacent pens, respectively. The infection force via this pathway is influenced by the accumulation of the virus in adjacent pens, factoring in the daily decay of the viral load ($\epsilon_1$)and the cleaning rate of fecal matter ($\epsilon_2$) from these pens. Transmission depends on the transfer of fecal matter from neighboring pens and the subsequent ingestion by pigs ($Q_i$). $N_{p,i-1}(t)$, $N_{p,i+1}(t)$, $I_{p,i-1}(t)$, and $I_{p,i+1}(t)$ denote the total number of pigs and the number of infected pigs in pen $i-1$ and pen $i+1$ at time $t$, for the left and right neighbors, respectively.

##### 2.2.3 Room-level air-borne transmission

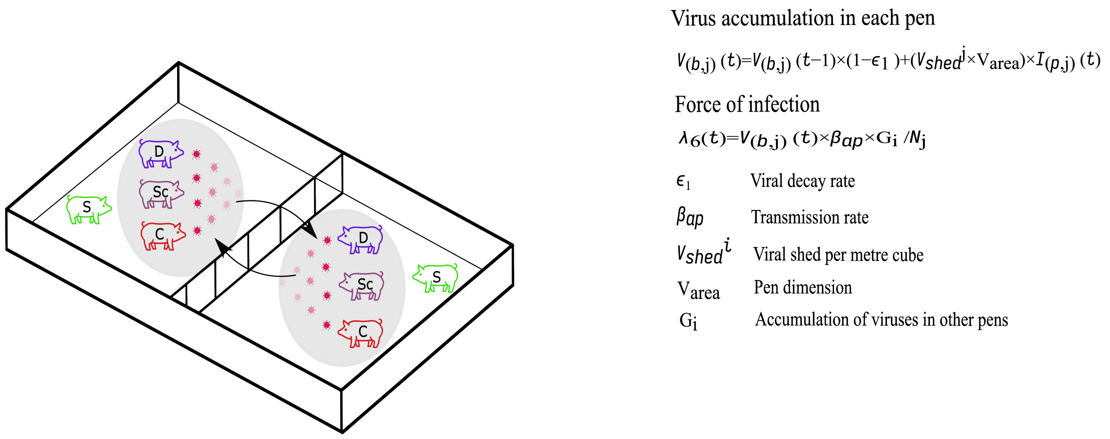


Figure 6: The schematic illustrates aerosol transmission between pens. The cumulative viral load is determined by the viral loads in all pens except the one being considered, with daily viral decay taken into account. A pen-to-pen network is constructed using a directed graph represented as $G = (V, E)$, where pens are defined as a set of vertices $V = v_1, v_2, ..., v_n$, and their directed edges $E$ signify the potential pathways between pens. The edge $(v_i, v_j)$ indicates the amount of airborne pathogens transmitted from pen $j$ to pen $i$. In this airborne transmission route, each pen $v_i$ is connected to every other pen $v_j$ where $i \neq j$. The viral load depends on the number of particles shed by an infectious pig per day, defined by the parameter $V^i_{shed}$ per cubic meter of air. $V_{area}$ determines the level of viruses present in each pen. $G_i$ represents the cumulative virus transferred to pen $i$ from all other pens. $\beta_{ap}$ is the aerosol transmission rate, and $N_j$ is the number of pigs in the neighboring pens.

#### 2.3 Human-mediated transmission at room level

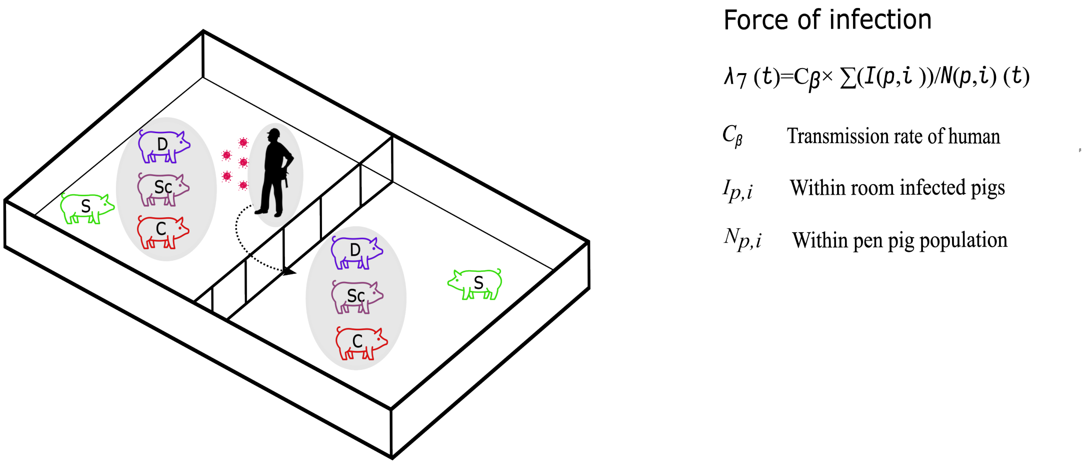


Figure 7: Figure depicting the human-mediated transmission. This route depends upon the total number of infected pigs present within the room. $C_\beta$ is the human transmission rate, $N_p$ is the total number of pigs in the room, and $I_p$ is the total number of infected pigs in the room.

,

### 3 Pen-to-pen pigs transferring sensitivity analyses

We investigate the dynamics of ASFV spread and the contribution of individual transmission routes to disease spread in rooms comprising 24 pens and 40 pens, under three different probabilities of transferring pigs (A) 5 %(base model), (B) 25 %, and (C) 50 %, respectively.

#### 3.1 Results for a room with 24 pens

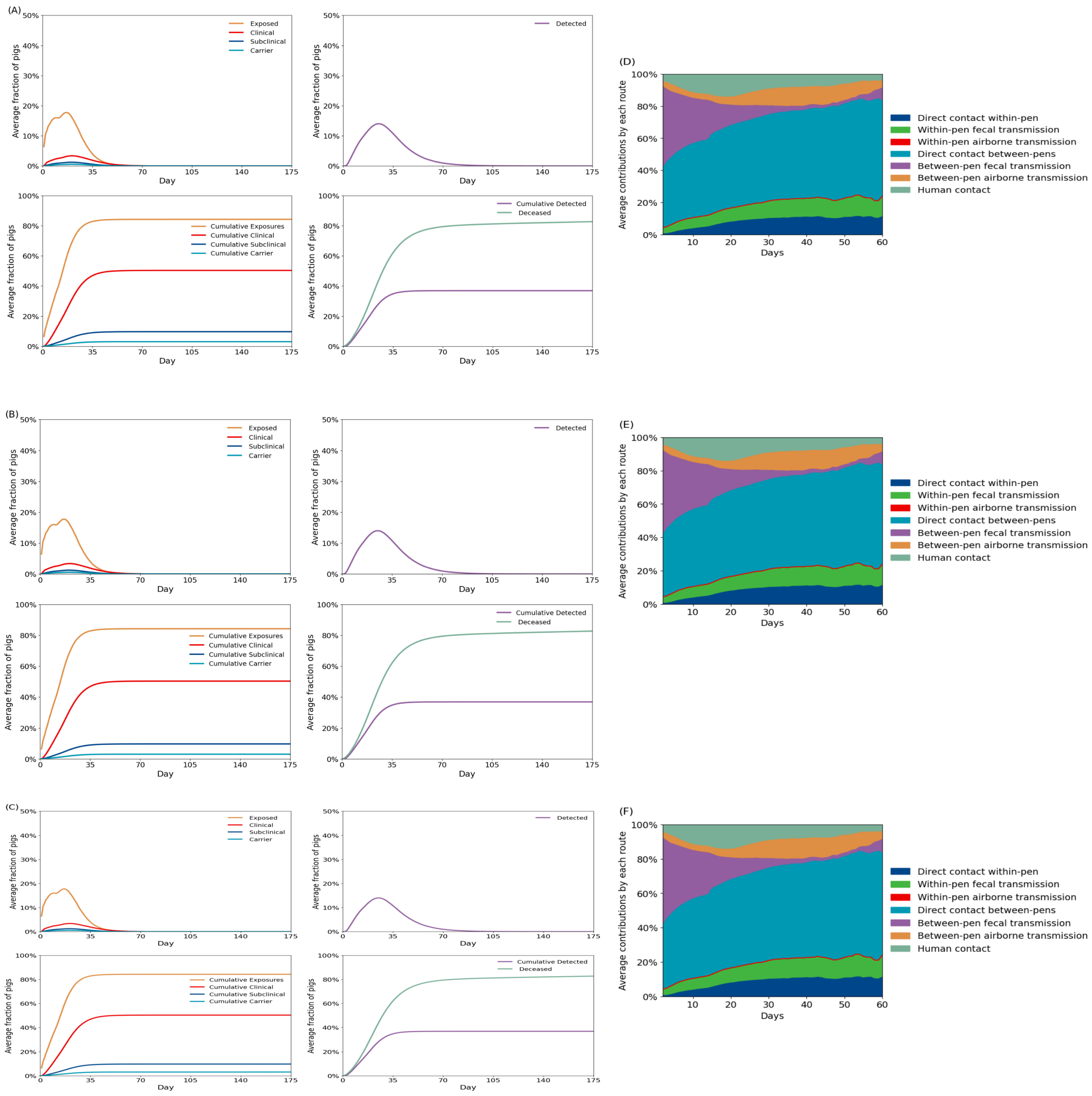


Figure 8: Disease dynamics for different rates of transfer of pigs for a room with 24 pens. The probability of transferring a pig on a day is fixed at 5 %(base model), 25 %, and 50 %, respectively.

#### 3.2 Results for a room with 40 pens

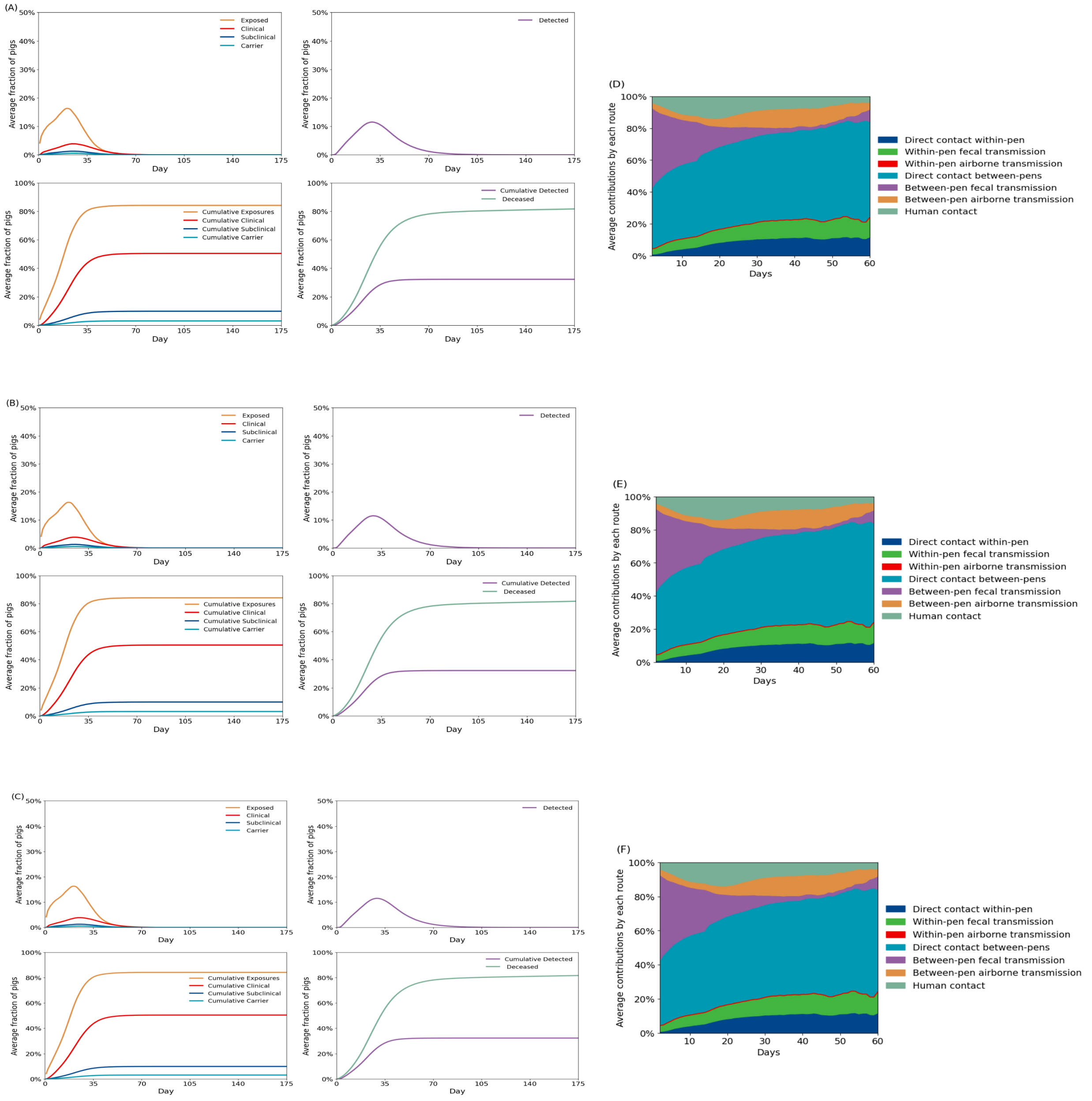


Figure 9: ASFV spread and pathways contribution (bottom panel) in a room with 40 pens (top panel), categorized by the daily probability of pig transfers: 5% (left), 25% (middle), and 50% (right).

### 4 Human-mediated transmission sensitivity analyses

We investigate the dynamics of ASFV spread and the contribution of individual transmission routes to disease spread in rooms comprising 40 pens, under varying human-mediated transmission rates.

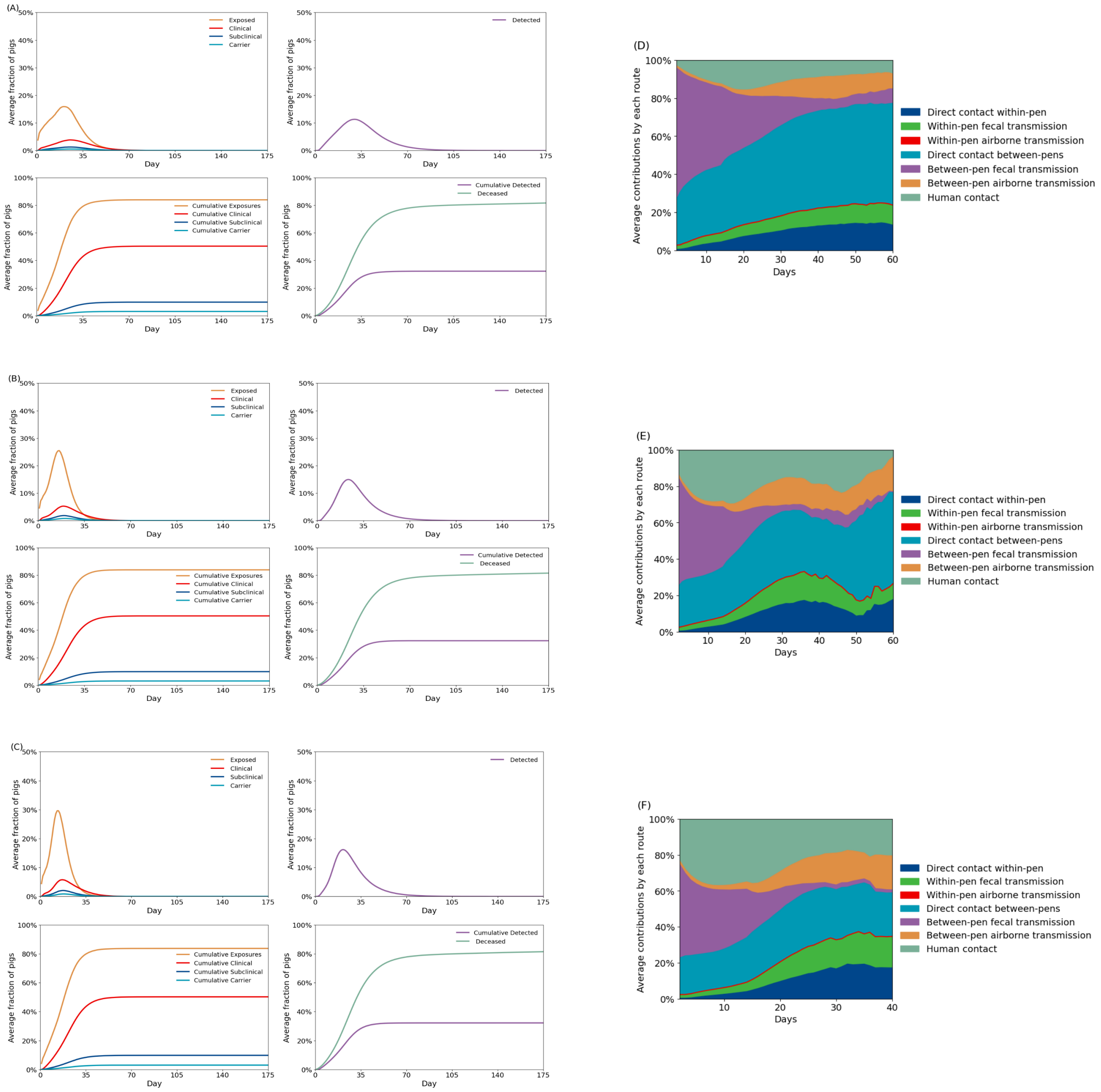


Figure 10: We investigate the spread of ASFV and the contribution of individual routes in a room consisting of 40 pens under different human-mediated transmission scenarios: A) and D) for the base model with $C_\beta$=0.01, B) and E) for $C_\beta$ =0.05, and C) and F) for $C_\beta$=0.1.

### 5 Mortality rate sensitivity analyses

We investigate the dynamics of ASFV spread and the contribution of individual transmission routes to disease spread in rooms comprising 40 pens, under varying mortality rates.

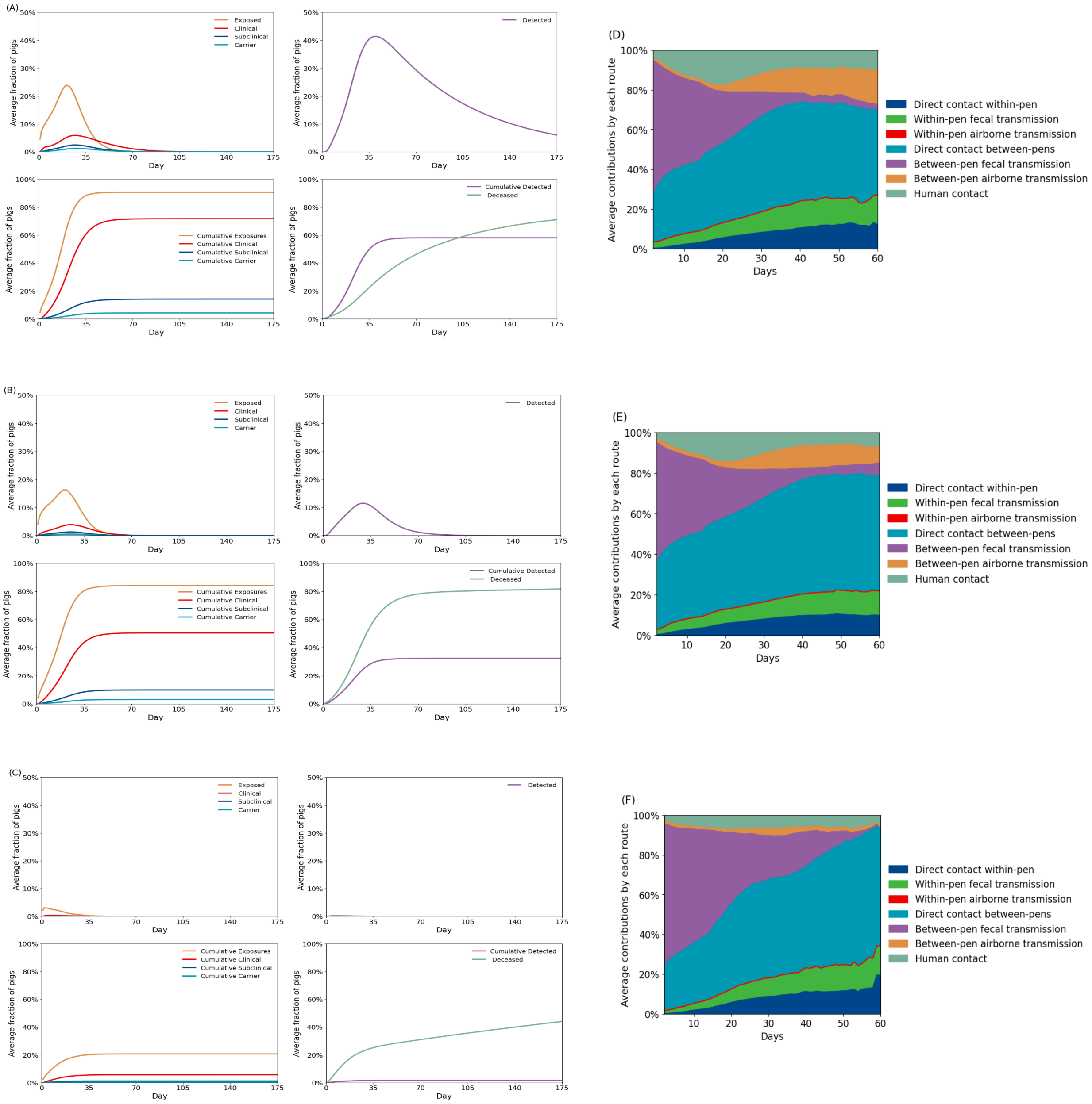


Figure 11: We investigate the spread of ASFV and the contribution of individual routes in a room consisting of 40 pens under different mortality rates: A) and D) for the low mortality rate where the mortality rate is reduced by five times, B) and E) for base mortality rate, and C) and F) for the high mortality rate where the mortality rate is increased by five times

### 6 Viral load dynamics

#### 6.1 Viral load across a room consisting of 40 pens over time

In this subsection, we investigate the viral load in each pen across three different time periods, considering both fecal and aerosol transmission routes. We subdivide the barn into three sections: Section I represents the pens closer to the fans, Section II represents the middle of the barn, and Section III comprises the pens closer to the exhaust fans. Our objective is to determine where the highest viral loads are located during the three different periods of ASFV spread.

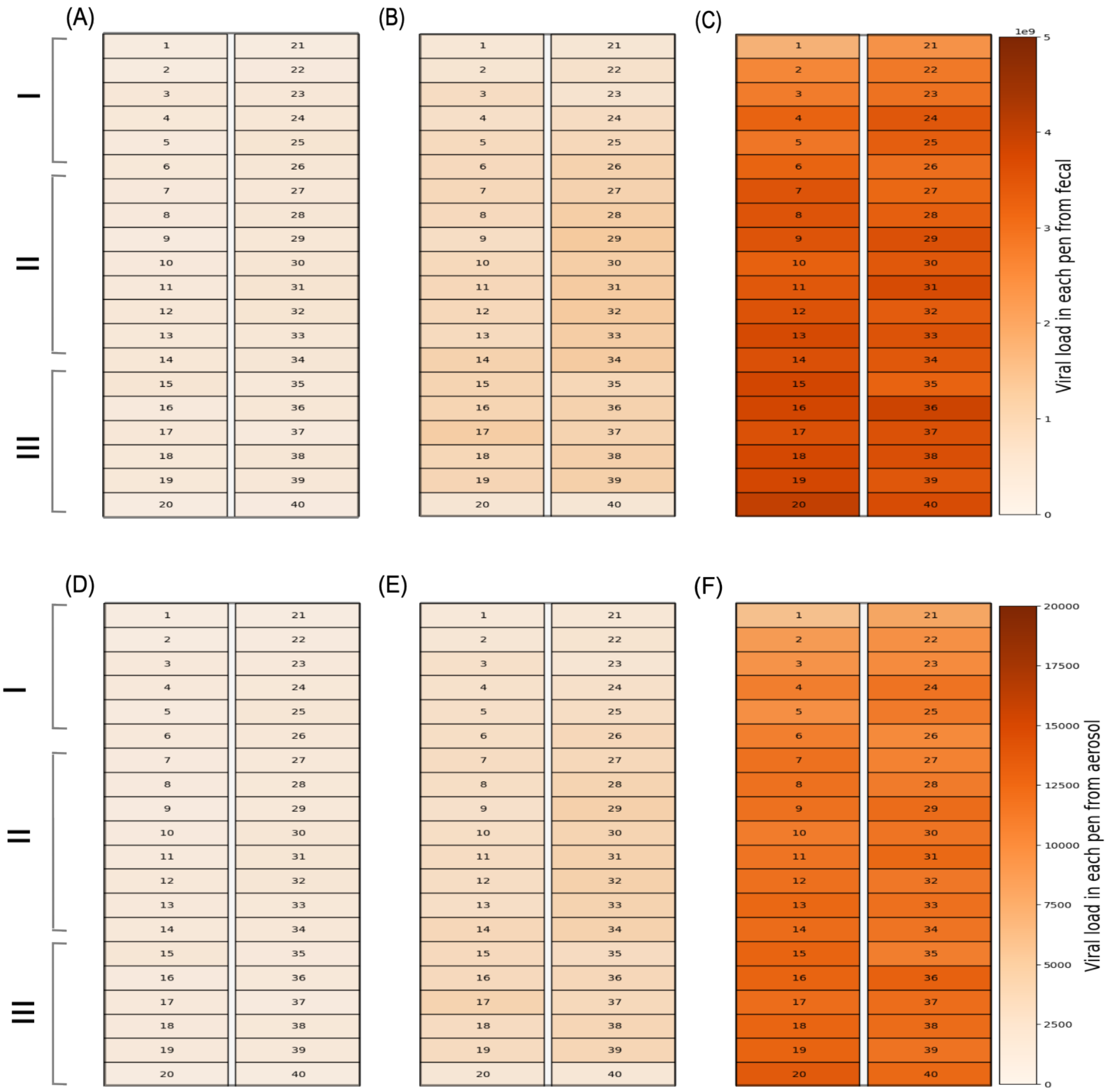


Figure 12: Average viral loads generated through fecal transmission (top panel) and aerosol transmission (bottom panel) across 40 pens over periods of (A, D) 5, (B, E) 10, and (C, F) 30 days.

#### 6.2 Average viral load across all rooms from day 1 to day 175 of ASFV spread

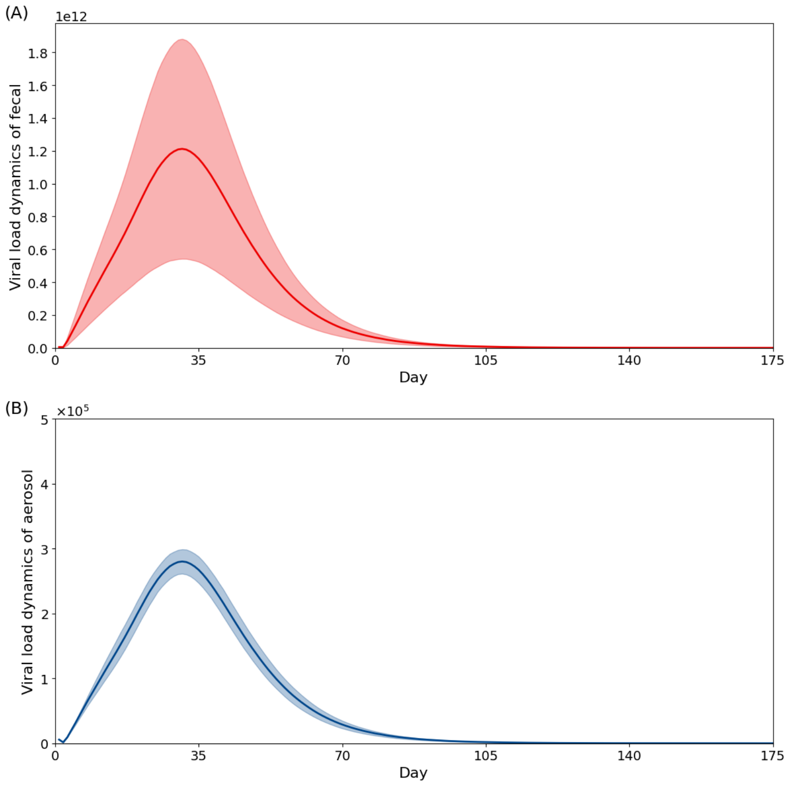


Figure 13: The average amount of viral loads generated through (A) fecal transmission and (B) aerosol transmission over 175 days of simulated outbreaks across all rooms. The vertical axis is the mean virus particles generated across all rooms. The solid line represents the mean viral load generated, and the shaded region represents the standard deviation at duration $t$(days).

### 7 ASFV dissemination at pen-level in rooms with 28 and 32 pens

In this subsection, we investigate the number of exposed, clinical, sub-clinical, and carrier pigs in each pen across three different time periods in a room comprising 32 pens. Section I represents the pens closer to the fans, Section II represents the middle of the barn, and Section III comprises the pens closer to the exhaust fans.

#### 7.1 Exposed pigs

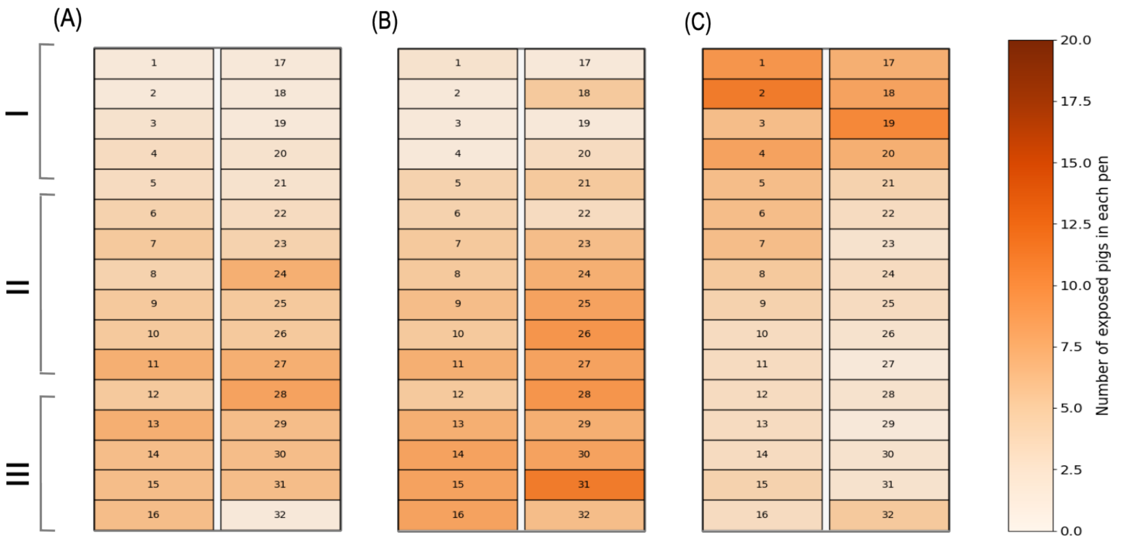


Figure 14: Number of exposed pigs in each pen within a room containing 32 pens over a period of (A) 5 days, (B) 10 days, and (C) 30 days.

#### 7.2 Clinical pigs

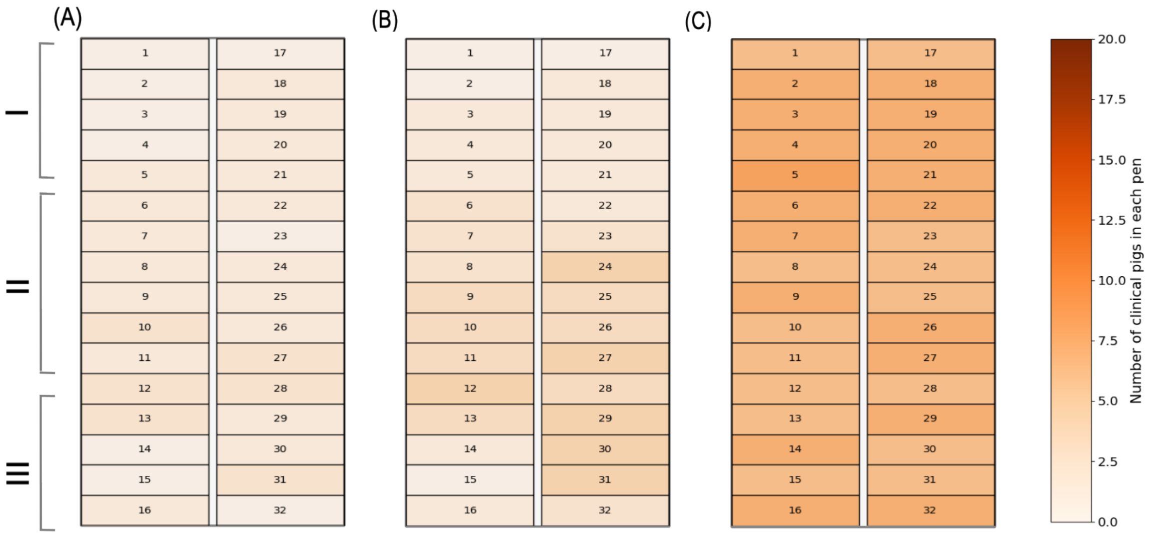


Figure 15: Number of clinical pigs in each pen within a room containing 28 pens over a period of (A) 5 days, (B) 10 days, and (C) 30 days.

##### 7.3 Sub-clinical pigs

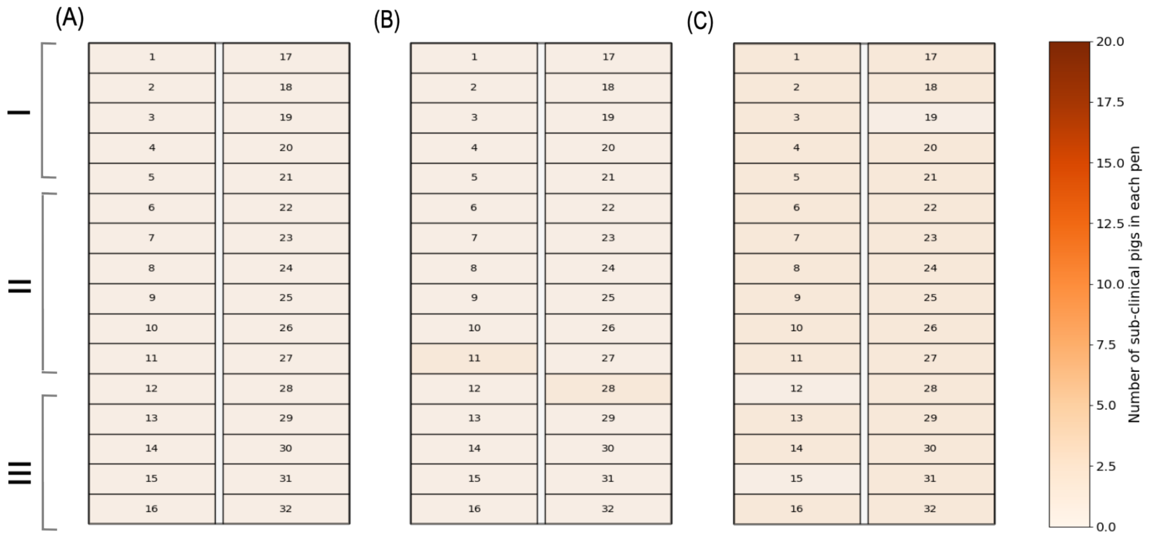


Figure 16: Number of sub-clinical pigs in each pen within a room containing 28 pens over a period of (A) 5 days, (B) 10 days, and (C) 30 days.

##### 7.4 Carrier pigs

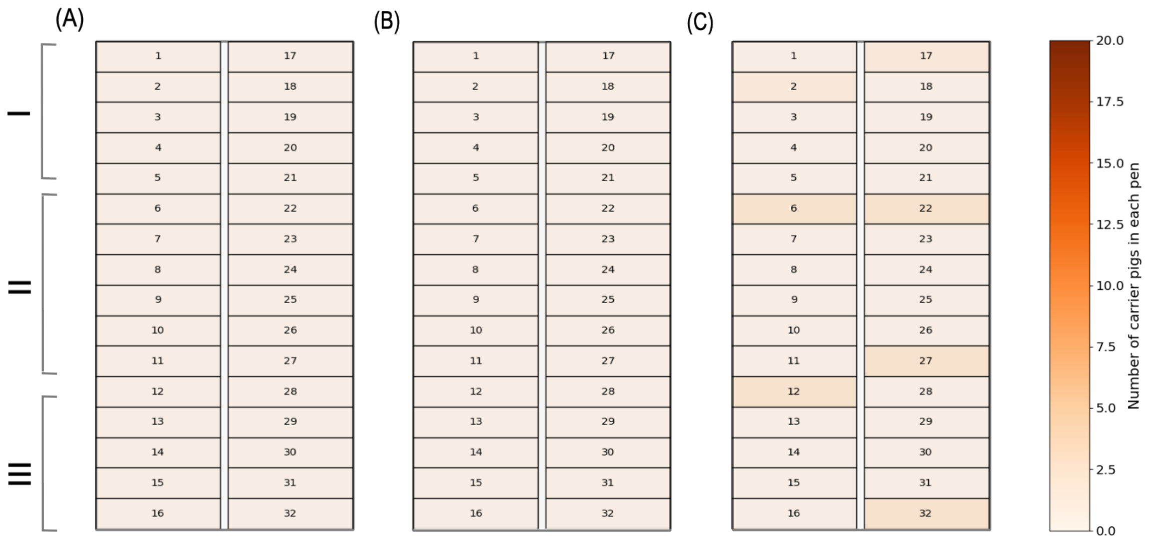


Figure 17: Number of carrier pigs in each pen within a room containing 28 pens over a period of (A) 5 days, (B) 10 days, and (C) 30 days.

### 8 Recovered and susceptible compartments dynamics over time

In this subsection, we present the average number of pigs that remain susceptible and those that have recovered over a period of 175 days of ASFV spread, observed across 7704 rooms.

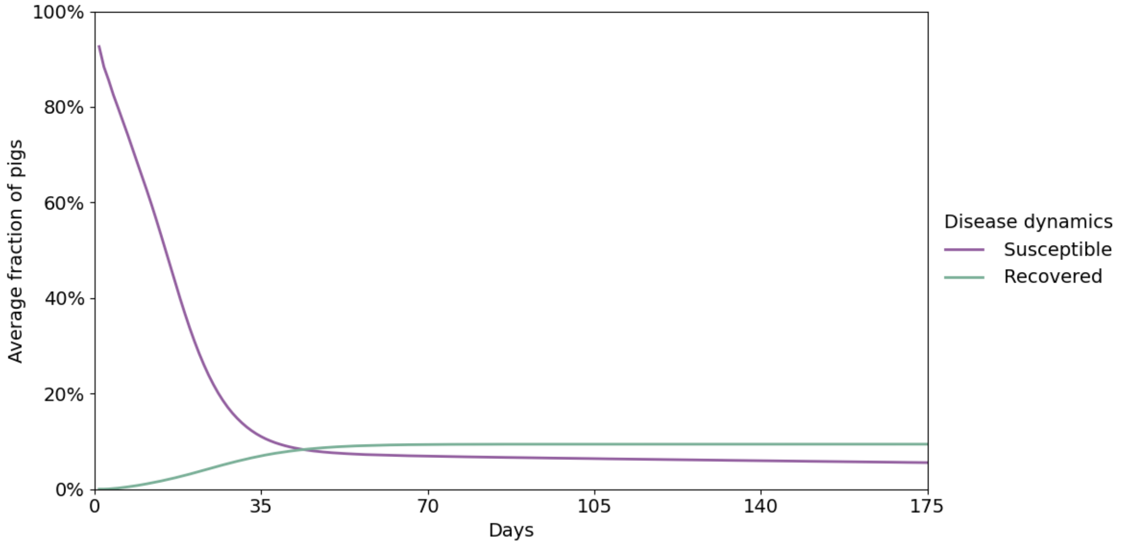


Figure 18: Average prevalence of pigs that are susceptible and recovered across all rooms over 175 days.
